# CD226 identifies effector CD8^+^ T cells during tuberculosis and costimulates recognition of *Mycobacterium tuberculosis*-infected macrophages

**DOI:** 10.1101/2025.01.22.634303

**Authors:** Tomoyo Shinkawa, Evelyn Chang, Tasfia Rakib, Kelly Cavallo, Rocky Lai, Samuel M. Behar

**Author notes:** Corresponding author: Samuel M. Behar, E-mail address (SMB).

## Abstract

CD8^+^ T cells defend against *Mycobacterium tuberculosis* (Mtb) infection but variably recognize Mtb-infected macrophages. To define how the diversity of lung parenchymal CD8^+^ T cells changes during chronic infection, cells from C57BL/6J mice infected for 6- and 41-weeks were analyzed by scRNA-seq. We identified an effector lineage, including a cluster that expresses high levels of cytotoxic effectors and cytokines, and dysfunctional lineage that transcriptionally resembles exhausted T cells. The most significant differentially expressed gene between two distinct CD8^+^ T cell lineages is CD226. Mtb-infected IFNγ-eYFP reporter mice revealed IFNγ production is enriched in CD226^+^CD8^+^ T cells, confirming these as functional T cells in vivo. Purified CD226^+^ but not CD226^−^ CD8^+^ T cells recognize Mtb-infected macrophages, and CD226 blockade inhibits IFNγ and granzyme B production. Thus, CD226 costimulation is required for efficient CD8^+^ T cell recognition of Mtb-infected macrophages, and its expression identifies CD8^+^ T cells that recognize Mtb-infected macrophages.

**One Sentence Summary:** Shinkawa et al. discover that CD226 is a functional marker that distinguishes effector from dysfunctional CD8^+^ T cells in the *Mycobacterium tuberculosis* (Mtb)-infected lung and has a crucial role in costimulating CD8^+^ T cell recognition of Mtb-infected macrophages.

## Introduction

*Mycobacterium tuberculosis* (Mtb), the bacterium that causes the human disease tuberculosis (TB), is an extraordinarily successful intracellular pathogen that coevolved with humans and developed ways to evade our host defenses. It is estimated that Mtb has infected 25% of the world’s population (*1*). Only 5-10% of infected people develop TB, which attests to the effectiveness of human immunity. Mtb-specific T cells are essential to prevent TB in humans and control infection in animal models (*2–4*). Mtb elicits strong CD8^+^ T cell responses associated with Mtb control in humans and experimentally infected animals (*4–16*). In the mouse TB model, polyclonal CD8^+^ T cells elicited by infection can restrict Mtb in vitro (*17*) and in vivo (*13*), and vaccine-elicited CD8^+^ T cell responses improve control of pulmonary TB (*18, 19*). CD8^+^ T cells are essential for the long-term survival of mice following intravenous Mtb infection (*12, 20*). These data show the potential of CD8^+^ T cell responses to combat Mtb infection. However, CD8^+^ T cell depletion leads to only modest reductions in survival after low-dose aerosol infection (*4*). Thus, the mechanisms of how CD8^+^ T cells mediate immunity, and how Mtb evades effective CD8^+^ T cell responses are poorly understood.

A priori, three scenarios could limit immunity mediated by CD8^+^ T cells. First, Mtb could have evolved to avoid CD8^+^ T cell recognition by producing decoy antigens or avoiding antigen entry into the class I MHC pathway. We described the inefficient presentation of the immunodominant TB10.4 antigen by Mtb-infected macrophages to CD8^+^ T cells (*21–23*) and found that only 10-15% of CD8^+^ T cells from the lungs of Mtb-infected C57BL/6J mice recognize infected macrophages (*24*). Second, CD8^+^ T cells might fail to express effector functions associated with Mtb control. Among human T cells, tri-cytotoxic cells that express perforin, granzymes and granulysin are associated with anti-mycobacterial activity (*7, 8, 10*). While the absence of a granulysin ortholog in the *Mus* genome could explain why mice are unable to clear Mtb, a suitable model to test this hypothesis is not currently available (*25*). Finally, Mtb-specific CD8^+^ T cells might become dysfunctional because of persistent stimulation, as has been observed during chronic viral infection and cancer. Continued T cell stimulation by persistent antigen induces a CD8^+^ T cell state known as exhaustion (*26, 27*). Exhausted CD8^+^ T cells have sustained expression of inhibitory receptors, reduced effector function, and diminished proliferation. Sustained inhibitory receptor expression, including PD-1 and TIM-3, on a subset of CD8^+^ T cells occurs in the murine TB model when the immune system is perturbed (*17, 28*) and in chronically infected mice (*29*). However, the underlying heterogeneity in phenotype and functionality of CD8^+^ T cells throughout infection is ill-defined. Whether T cell exhaustion occurs in human and non-human primates is also less clear (*30–32*).

To investigate the factors that compromise CD8^+^ T cell immunity, we used scRNA-seq and paired scTCR-Seq to define the heterogeneity of lung parenchymal CD8^+^ T cell responses during Mtb infection, early after the establishment of T cell-mediated control in infected C57BL/6J mice (i.e., 6 weeks post-infection, wpi) and during chronic infection (i.e., 41 wpi). CD8^+^ T cell responses are diverse and evolve over time. We find two major lineages of differentiation. One is an effector lineage, including a cluster of polyfunctional effectors; the other is a dysfunctional lineage that are transcriptionally similar to exhausted CD8^+^ T cells described during chronic LCMV infection and cancer. Dysfunctional CD8^+^ T cells increase in proportion over time. Notably, the gene encoding CD226 (DNAX accessory molecule-1; DNAM-1) is the most significantly differentially expressed gene (DEG) between effector and dysfunctional CD8^+^ T cell lineages. CD226 was first discovered as an adhesion molecule that enhances cytotoxicity in NK and T cells (*33*). Recent studies show that CD226 expression by CD8^+^ T cells is a favorable prognostic factor for cancer outcomes in humans and mice, and the efficacy of immune checkpoint blockade requires CD226 expression on CD8^+^ T cells (*34–37*). Using flow cytometry, IFNγ reporter mice, and an in vitro Mtb infection model, we show that CD226 is a marker that identifies effector CD8^+^ T cells with retained polyfunctionality, which persists during chronic infection in vivo. In vitro, not only does CD226 identify CD8^+^ T cells that recognize Mtb-infected macrophages, but blocking CD226 impairs the production of effector molecules, including IFNγ and granzyme B by CD8^+^ T cells, suggesting that CD226 is required for efficient recognition of infected macrophages. Dysfunctional CD8^+^ T cells, characterized by a lack of CD226 expression and inability to recognize infected macrophages, emerge as early as 4 wpi and increase massively in proportion over time. Our data provide novel mechanistic insights into effective CD8^+^ T cell immune response and its failure during Mtb infection.

## RESULTS

### Nine CD8^+^ T cell states are identified after Mtb infection

To assess the heterogeneity of CD8^+^ T cells both early and late during chronic Mtb infection, we generated single cell (sc) RNA-seq and TCR-seq datasets of lung parenchymal (CD45-IV^−^) CD3^+^ T cells, 6- and 41-weeks post-infection (wpi) (Fig.S1A). After quality control, cells expressing both *CD8a* and *CD8b1* were selected for analysis (n=9,870). To avoid T cell receptor (TCR) genes influencing clustering, we removed TCR genes before scaling and dimensionality reduction (*38*). Unsupervised clustering identified nine clusters of CD8^+^ T cells (Fig.1A, B, Table S1). Cluster designations were based on immunological signatures and key differentially expressed genes (Fig.1C, D, Table S1).

**Figure 1.**
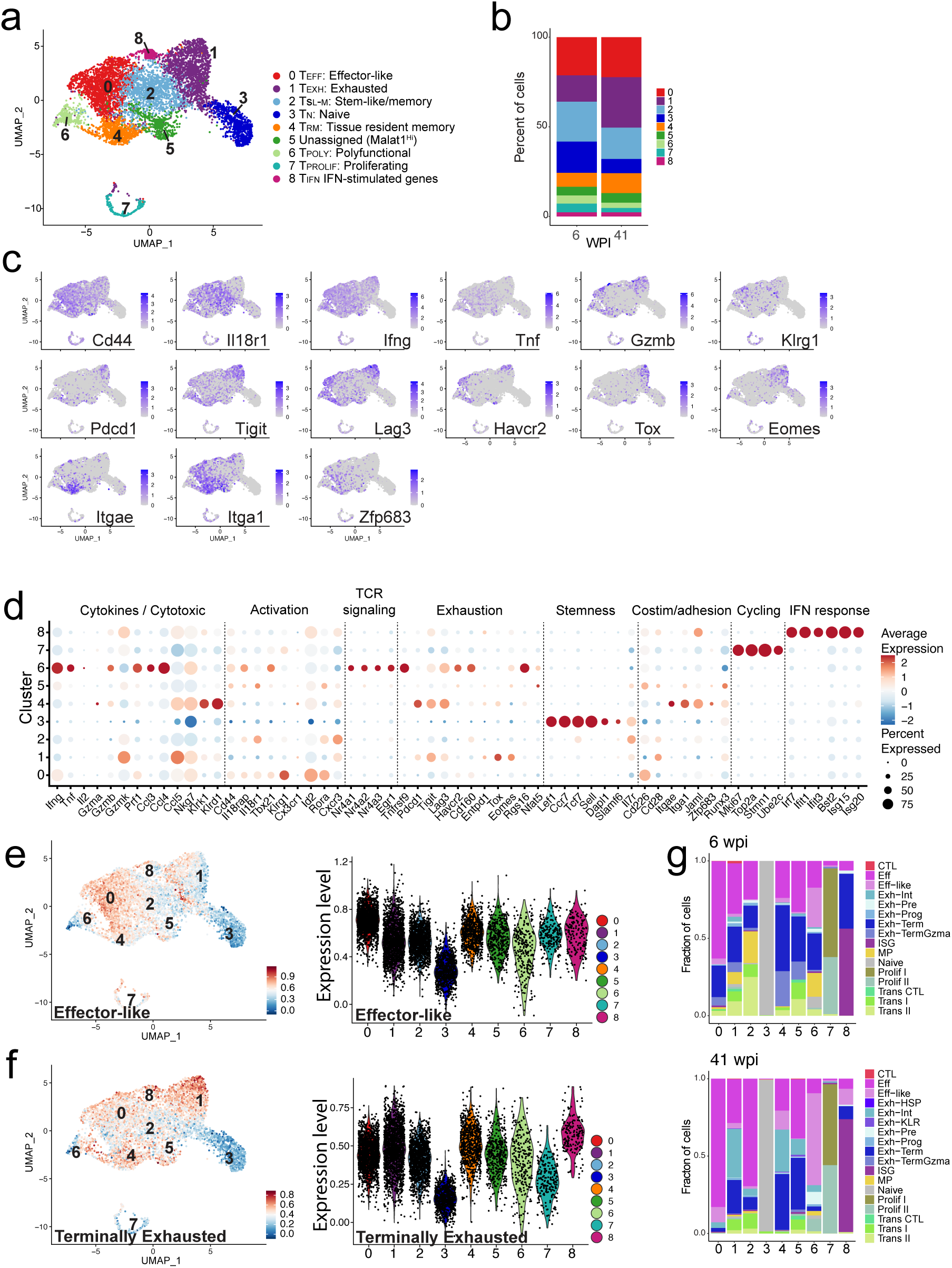
Nine distinct lung parenchymal CD8^+^ T cell states are identified during Mtb infection. **A)** UMAP visualization of scRNAseq data of lung parenchymal CD8^+^ T cells from mice infected with Mtb for 6 and 41 weeks. **B)** Stacked bar graphs depict the cluster distribution at 6 and 41 wpi. **C)** Feature plots of the indicated gene transcripts. **D)** Dot plot of selected function-associated genes among the CD8^+^ T cell clusters. **E, F)** UMAP colored based on enrichment of ‘effector-like’ or ‘terminally exhausted’ gp33-specific CD8^+^ T cell signatures from LCMV Clone 13 infection [GSE122712](*39*) and violin plots of the enrichment score. **G)** Transferred cluster annotations of gp33-specific splenic CD8^+^ T cells from the combined datasets of LCMV-Armstrong- and LCMV-Cl13-infected mice [GSE199565](*58*) to our CD8^+^ T cell clusters at 6 and 41 wpi.

Cluster 0 is designated as effector-like T cells (hereafter referred to as ‘TEFF/0’) based on the expression of *Klrg1*, *Cd226*, *Id2, Rora,* and *Ifng*. A large percentage of cells are TEFF/0, even at 41 wpi. CD8^+^ T cells in Cluster 1 express high levels of co-inhibitory receptors *Pdcd1*, *Tigit*, *Lag3,* and *Gzmk* and transcription factors (TFs) associated with T cell exhaustion (*Tox* and *Eomes*) (Fig.1A, C, D, Table S1). CD8^+^ T cells in Cluster 1 have also lower expression of adhesion and costimulatory molecules, including *Cd226* and *Jaml*, and cytokines, including *Ifng* and *Tnf*. Therefore, Cluster 1 is defined as exhausted (TEXH/1) based on their similarity to exhausted CD8^+^ T cells during LCMV clone 13 infection and in tumors (*39*). Downregulation of *Il18r1* on TEXH/1 suggests they lose responsiveness to IL-18 (*40*). TEXH/1 is the largest population at 41 wpi (Fig.1B). CD8^+^ T cells in Cluster 2 express *Il7r*, *Tcf7*, *Il18r1*, and *Cxcr3*. These cells are designated as stem-like or memory CD8^+^ T cells (TSL-M/2) as they express few, if any, genes associated with effector function. The CD8^+^ T cells in Cluster 3 are designated as naïve (TN/3) based on their expression of *Ccr7*, *Sell, Lef1, Dapl1*, *and Tcf7* (Fig.1D, Table S1). Cluster 4 CD8^+^ T cells express the adhesion molecules *Itgae* (CD103) and *Itga1* (CD49a) consistent with tissue-resident memory T cells (TRM/4) (Fig.1A-D). Their expression of *Gzmb, Klrk1* (NKG2D), and *Klrd1* (CD94) indicates potential cytotoxic T cell (CTL) activity. Interestingly, expression of *Pdcd1, Tigit*, *and Lag3* in TRM/4 suggests persistent TCR stimulation (*41*). Enrichment of TGFβ signaling signature (*42*) in TRM/4 is consistent with TGFβ affecting differentiation and maintenance of TRM (*43–47*) and co-inhibitory receptor expression (*48*) (Fig.S1B).

The remaining 25% of CD8^+^ T cells are distributed among Clusters 5-8. Cluster 5 shares features with activated CD8^+^ T cells, including upregulation of components of AP-1 family TFs, *Fosb*, and *Jun*. However, they express mitochondrial and nuclear genes, including *Malat1* (*49*) and had low recovery of paired TCRα and TCRβ transcripts, suggesting dying or low-quality cells. Cluster 6 is designated as polyfunctional CD8^+^ T cells (TPOLY/6) based on their high expression of *Ifng, Tnf, Prf1*, *Gzmb, Ccl3, and Ccl4*. Expression of Nr4a family TFs and *Tnfrsf9* suggests active TCR signaling. The expression of exhaustion-related genes, including *Havcr2, Lag3,* and *Rgs16,* could indicate chronic TCR stimulation (*50*). Cluster 7 CD8^+^ T cells are proliferating (T_PROLIF_/7) based on *Mki67, Top2a, Stmn1*, and *Ube2c* expression. Finally, CD8^+^ T cells in Cluster 8 have a type I interferon (IFN I) signature (TIFN/8) based on *Ifit1, Ifit3, Bst2, Isg15*, and *Isg20* expression.

To explore TF regulon activity in each CD8^+^ T cell state, we performed Single-cell Regulatory Network Inference and Clustering (SCENIC) (*51*). We identified increased expression of target genes and TF activity of Rorα and Klf6 in TEFF/0 (Fig.S1C). TEXH/1 displayed high Eomes activity, consistent with Eomes promoting CD8^+^ T cell exhaustion (*52*). TF activity of Irf1, −2, −7, and −9, and Stat1 and −2, are enriched in TIFN/8, consistent with IFN I responses (Fig.S1C). We inferred cytokine signaling activity using Cytokine Signaling Analyzer (CytoSig) (Fig.S1D)(*53*). A pronounced response to IL-18 is predicted for TEFF/0 and TSL-M/2, supporting their designation as effector and memory-like CD8^+^ T cells (*40*). TIFN/8 is expected to have an elevated response to interferons and to IL-27 which induces co-inhibitory gene programs (*54*).

To confirm the clusters’ identity, we applied gene signatures from a dataset of CD8^+^ T cells induced by Lymphocytic Choriomeningitis Virus (LCMV) Clone 13 infection. The “Effector-like signature” has the highest score in TEFF/0 and the lowest score in TEXH/1 and TN/3 (Fig.1E)(*39*). The “Terminally exhausted signature” also from LCMV Clone 13 infection, has the highest score in TEXH/1 and TIFN/8 and the lowest score in TN/3 and T_PROLIF_/7 (Fig.1F)(*39*). The high “Terminally exhausted signature” score for TIFN/8 supports IFN I driving CD8^+^ T cell dysfunction (*55, 56*). To further substantiate our cluster annotations, we performed a label transfer of other dataset which include CD8^+^ T cells from both acute and chronic LCMV infection to our dataset (*57, 58*). Nearly all TEFF/0 cells are identified as effector (TEFF), a CD8^+^ T cell state observed exclusively after acute LCMV infection, but not chronic LCMV infection (Fig.1G, Fig.S1E, F). CD8^+^ T cells from chronic LCMV infection (e.g. “Exh−Int,” “Exh−term,” Fig.S1F) project onto TEXH/1 and TRM/4, which increase in proportion at 41 wpi. This analysis confirms that TEFF/0 and TPOLY/6 are in effector states, while TEXH/1 is in an exhausted state. TRM/4 has a mixed feature of effector and exhausted states. In summary, we identify nine distinct clusters of CD8^+^ T cells, present both early and late after Mtb infection, including clusters with multiple effector molecule expression and clusters that resemble exhausted CD8^+^ T cells.

### *Cd226* distinguishes between effector-like and dysfunctional CD8^+^ T cell lineages

To understand lineage relationship of CD8^+^ T cell clusters and to infer which responses are driven by Mtb infection, we leveraged paired scTCR-seq analysis (Table S2). Clonal expansions are a cardinal feature of T cell responses and cells with identical TCR rearrangements indicate a common origin. T cells with identical CDR3α and CDR3β nucleotide sequences were defined as a clonotype. We identified 1,187 unique TCRαβ clonotypes among 5,833 cells. Dramatic expansions were detected in TEFF/0 consistent with an effector CD8^+^ T cell response to Mtb infection (Fig.2A). In contrast, expansions of clonotypes in TEXH/1 were smaller and most clonotypes in TN/3 were unique, consistent with their exhausted and naïve T cell designation, respectively.

**Figure 2.**
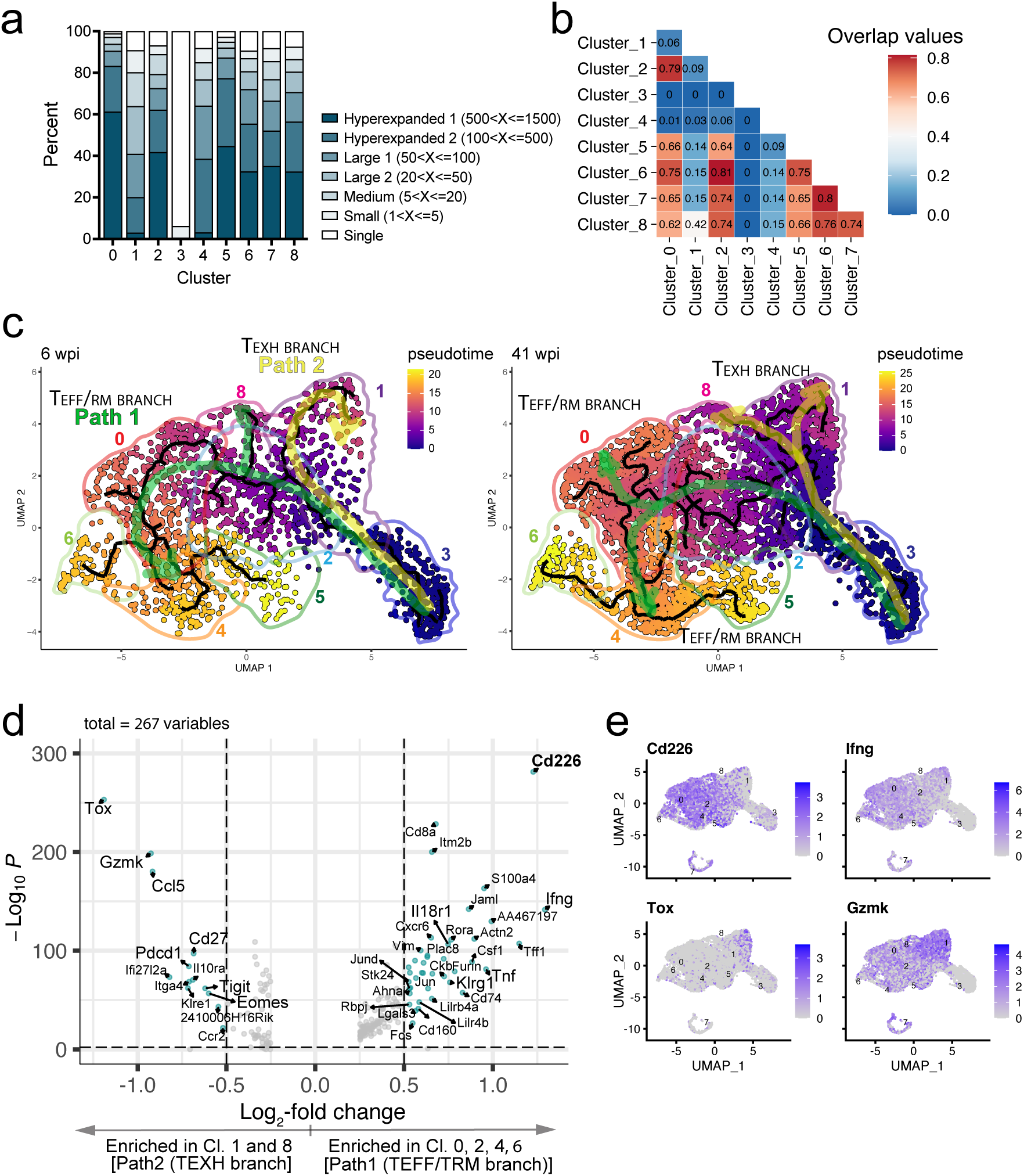
Developmental bifurcation of effector-like/tissue-resident memory and exhausted CD8^+^ T cells differentiated by *Cd226* expression. **A)** Stacked bar plots of clonal sizes among clusters. **B)** Plot denoting the Morisita index value for each cluster comparison demonstrating the TCR clonotype overlap. **C)** Single-cell trajectories constructed using Monocle3. UMAP shows cells colored by pseudotime values along the trajectory (black line) at 6 wpi (left) and 41 wpi (right). Cluster number is denoted on the UMAP. T_PROLIF_/7 is excluded from the analysis. **(D)** Volcano plot of DEGs between TEFF/RM branch [TEFF/0, TSL-M/2, TRM/4, and TPOLY/6] and TEXH branch [TEXH/1 and TIFN/8]. Genes with log2FC >0.5 and an adjusted p value < 0.01 are designated by a green filled circle. (**E)** Feature plots of *Cd226*, *Ifng*, *Tox*, and *Gzmk* transcripts.

The Morisita index was used to quantify TCR sharing between clusters in the combined 6 and 41 wpi dataset (Fig.2B). There was significant clonotype sharing between TEFF/0 and TSL-M/2 or TPOLY/6, and little or no sharing with TEXH/1. The relation between the clusters was independently assessed by pseudotime analysis using Monocle3 (*59, 60*) with TN/3 as the root cells. Path 1 led to functional branch, including TEFF/0, TSL-M/2, TRM/4, TPOLY/6 (hereafter, referred to as ‘TEFF/TRM branch’) while Path 2 led to TEXH/1 (hereafter, referred to as ‘TEXH branch’) (Fig.2C). Interestingly, TIFN/8 cells are derived from the TEFF/TRM branch at 6 wpi, but they are derived from the TEXH branch at 41 wpi (Fig.2C). TEFF/0, TRM/4, and TPOLY/6, cluster separately and distantly from TEXH/1, and have little TCR overlap. This is consistent with previous observations that T cell exhaustion represents a distinct T cell differentiation program from the effector differentiation program (*61, 62*). In contrast, CD8^+^ T cells in TSL-M/2 could give rise to TEFF/0 and TPOLY/6 based on TCR sharing and pseudotime analysis. Thus, CD8^+^ T cells follow at least two divergent differentiation trajectories with different functional fates.

We next sought to identify the most differentially expressed genes (DEGs) between the TEFF/TRM and the TEXH branch. We identified 267 DEGs (Fig.2D, Table S3). Cells on the TEXH branch are enriched for *Tox, Gzmk, Tigit, Pdcd1, Eomes, Il10ra, Ccl5,* and *Cd27.* Cells on the TEFF/TRM branch are enriched for *Cd226, Ifng, Tnf, Jaml, Klrg1,* and *Il18r1*. Remarkably, *Cd226* is the top DEG enriched in TEFF/TRM branch. Together with *Ifng*, *Cd226* distinguishes CD8^+^ T cells in TEFF/TRM branch from those expressing high levels of *Gzmk* and exhaustion-related genes, including *Tigit, Pdcd1, Tox*, and *Eomes* (Fig.2D, E).

### *Cd226* expression is associated with CD8^+^ T cell effector responses

To substantiate our finding that *Cd226* distinguishes between two distinct CD8^+^ T cell lineages (i.e., TEFF/TRM vs. TEXH), lung parenchymal CD226^+^ or CD226^−^ CD44^+^CD8^+^ T cells were purified from infected mice and analyzed by bulk RNAseq (Fig.S2A; Table S4). We observed 202 DEGs between CD226^+^ and CD226^−^ CD8^+^ T cells (Fig.3A). The CD226^+^CD8^+^ T cells expressed higher levels of genes associated with effector function (*Ifng*), terminal differentiation (*Klrg1*, *Havcr2*), and proliferation (*Mki67*, *Top2a*). Gene set enrichment analysis (GSEA) confirmed that CD226^+^CD8^+^ T cells were enriched in signatures related to effector CD8^+^ T cells (Fig.3B). In contrast, the CD226^−^ CD8^+^ T cells expressed genes associated with T cell exhaustion (*Pdcd1, Tox*). To further corroborate the correlation of *Cd226* expression with features of effector cells, we reanalyzed scRNA-Seq data using Monocle3 and identified 49 modules of coregulated DEGs (Table S5). Module 22 is enriched in the TEFF/TRM lineage (Fig.S2B), includes the GO terms “regulation of immune effector process,” “leukocyte migration,” “interferon−gamma production,” and “regulation of cytokine production involved in immune response” (Fig.S2C), and includes *Cd226*. Thus, *Cd226* expression identifies CD8^+^ T cells with effector program that undergo terminal differentiation, express high *Ifng*, and resist the expression of exhaustion-associated genes including *Tox* and *Pdcd1*.

**Figure 3.**
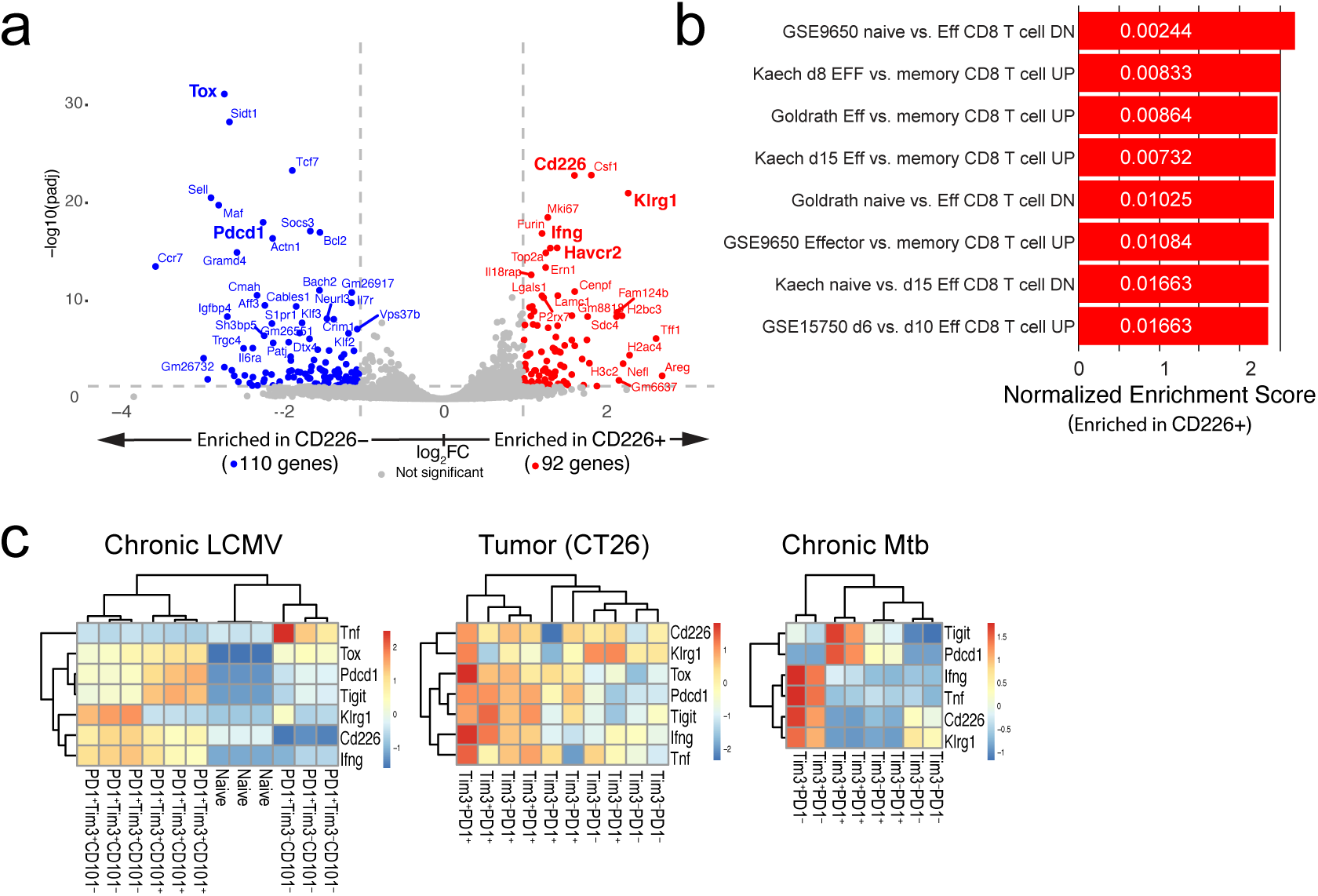
*Cd226* expression identifies CD8^+^ T cells with effector program during Mtb infection. **(A)** Volcano plot of DEGs between parenchymal CD226^+^ and CD226^−^ CD44^+^CD8^+^ T cells from the lungs of Mtb-infected mice at 24-28 wpi determined by RNAseq (log2FC>1, adjusted P value ≤ 0.05). **(B)** GSEA of differentially expressed genes in CD226^+^ CD8^+^ T cells versus CD226^−^ CD8^+^ T cells with log2FC>1 and adjusted P value ≤ 0.05 using MSigDB immunologic signature gene sets (C7). Adjusted P values are denoted in the bar. (**C)** Heatmaps of *Cd226*, *Ifng*, *Tnf, Pdcd1*, *Tigit*, *Klrg1,* and *Tox* expression (presented as z-score) for indicated CD8^+^ T cell subsets flow-sorted by PD-1 and TIM-3 (and CD101 in LCMV Clone13) expression determined by RNA-seq for LCMV Clone13(*63*) (left, n=3 donor), CT26 colon carcinoma(*64*) (middle, n=3-4 donor), and TB(*29*) (right, n=2 donor).

We next asked whether *Cd226* expression by CD8^+^ T cells is altered during other chronic diseases. CD8^+^ T cells in LCMV Clone 13 infection and tumor models exhibit distinct functional phenotypes defined by two co-inhibitory receptors: PD-1 and TIM-3. After LCMV Clone 13 infection, *Cd226* expression by CD8^+^ T cells is associated with a terminally exhausted PD-1^+^TIM-3^+^ phenotype (*63*)(Fig.3C). The results in the CT26 colon carcinoma model are not as clear, but *Cd226* expression is biased towards terminally exhausted PD-1^+^TIM-3^+^ CD8^+^ T cells (Fig.3C)(*64*). In contrast, reanalysis of our previous data finds that during TB, *Cd226* is mostly expressed by PD1^−^TIM3^+^ CD8^+^ T cells and not terminally exhausted PD-1^+^TIM-3^+^ T cells (*29*) (Fig.3C). Thus, the pattern of *Cd226* expression by CD8^+^ T cells with effector features but not exhausted CD8^+^ T cells during Mtb infection differs from its expression during chronic LCMV infection or cancer. Furthermore, PD1^−^TIM3^+^ CD8^+^ T cells, which are abundant during Mtb infection, are poorly described in chronic LCMV or tumor models (*64, 65*). In summary, *Cd226* expression identifies CD8^+^ T cells with an effector program during Mtb infection.

### CD226^+^CD8^+^ T cells express more IFNγ in vivo

As *Cd226* is highly expressed by CD8^+^ T cells with effector program at the RNA level, we next measured its cell surface expression by naïve splenic CD8^+^ T cells and lung CD8^+^ T cells isolated at 4, 11, 21, 30, and 43 wpi (Fig.4A, B, C). As previously reported, nearly all CD8^+^ and 20-40% of CD4^+^ splenic T cells from uninfected mice express CD226 by flow cytometry (Fig.4A, B). In the lungs of infected mice, 90-100% of parenchymal (CD45-IV^−^) antigen-experienced CD44^+^ CD62L^−^ CD8^+^ T cells initially express CD226; however, CD226 expression steadily declines during chronic infection (Fig.4A, B). After 40 wpi, ∼40% of parenchymal CD8^+^ T cells lack CD226 expression. In contrast, 50∼85% of CD4^+^ T cells express CD226, which fluctuates over the course of infection, but does not change significantly. Compared to naïve CD44^−^CD62L^+^ CD8^+^ T cells, antigen-experienced CD44^+^CD62L^−^ CD8^+^ T cells increase CD226 expression levels after infection (Fig.4C). CD226 is highly expressed by CD4^+^ and CD8^+^ T cells, but also γδ T and NK cells (Fig.S3A, B). Like in the lung, CD8^+^ T cells in the mediastinal lymph nodes (LN) and spleens of Mtb-infected mice gradually lose CD226 expression over time (Fig.S3C). Thus, CD226 is highly expressed by lung parenchymal CD8^+^ T cells, consistent with our scRNAseq data, and its expression decreases over time during Mtb infection.

**Figure 4.**
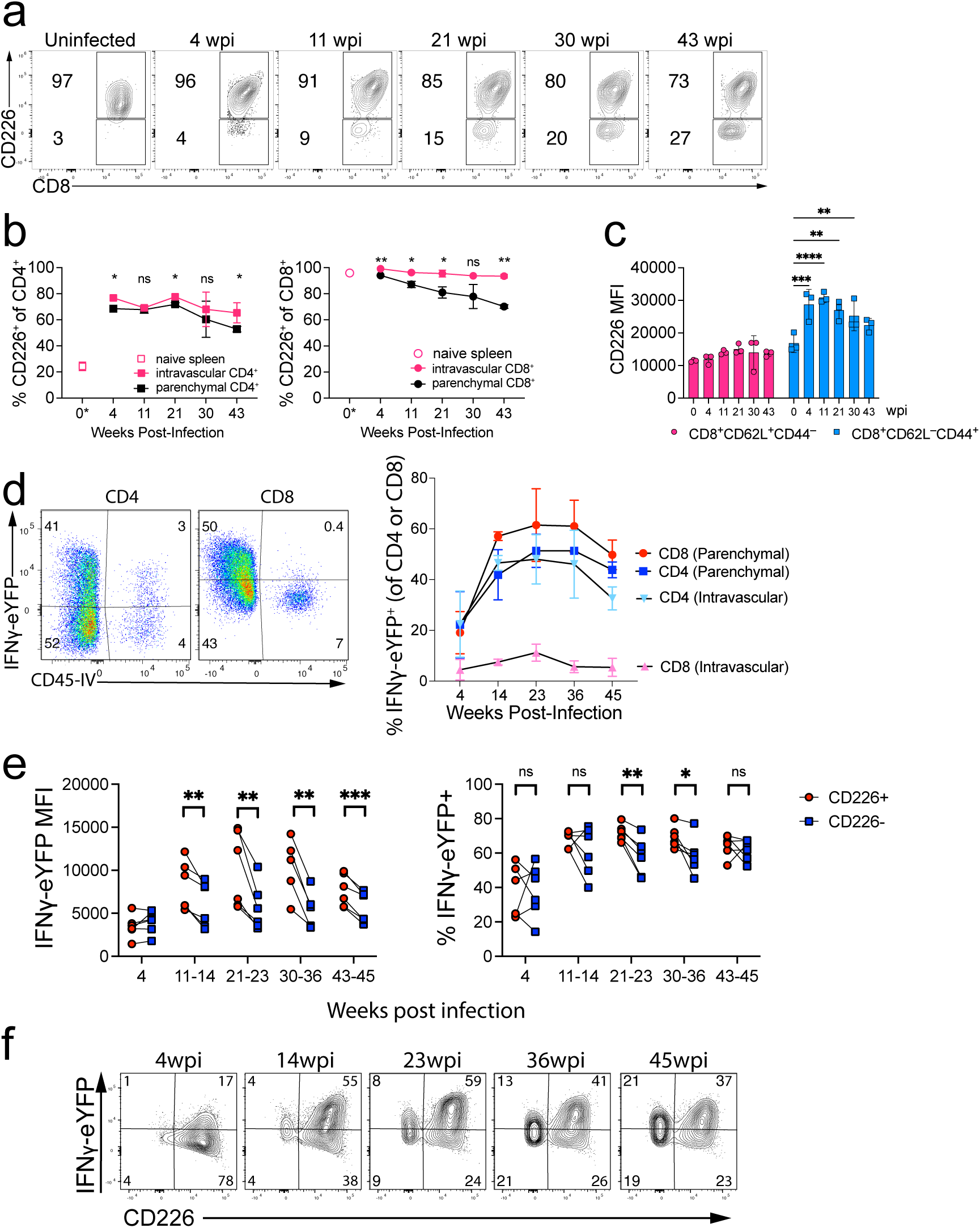
CD226^+^CD8^+^ T cells are activated in vivo during Mtb infection. (**A-C**) CD226 expression by CD8^+^ or CD4^+^ T cells from spleens of uninfected mice or CD8^+^ or CD4^+^ T cells from lungs of Mtb-infected mice was determined by flow cytometry. **(A)** Representative flow cytometry plots. **(B)** Percentage of CD226 expression by naïve splenic T cells from uninfected mice (0 wpi) or lung CD45-IV^+^ or CD45-IV^−^ T cells from Mtb-infected mice. CD4^+^ T cells (left) and CD8^+^ T cells (left) are shown. **(C)** Mean fluorescence intensity (MFI) of CD226 by CD8^+^ T cells from spleens of uninfected mice (0 wpi) or naïve (CD44^−^CD62L^+^) or antigen-experienced (CD44^+^CD62L^−^) lung CD45-IV^−^ CD8^+^ T cells. **(D)** IFNγ-eYFP expression by antigen-experienced (CD44^+^CD62L^−^) CD4^+^ or CD8^+^ T cells by lung CD45-IV^+^ or CD45-IV^−^ from Mtb-infected IFNγ-eYFP reporter mice. Shown are representative flow cytometry plots at 14 wpi (left) and quantification (right). **(E)** Quantification of IFNγ-eYFP MFI and percentage of lung antigen-experienced (CD44^+^CD62L^−^) CD226^+^ versus CD226^−^ CD8^+^ T cells from IFNγ-eYFP reporter mice infected with Mtb. **(F)** Representative flow cytometry plots of CD45-IV^−^ antigen-experienced (CD44^+^CD62L^−^) CD8^+^ T cells from Mtb-infected IFNγ-eYFP reporter mice for the indicated duration. **(A-F)** Data from three mice per group, representative of three experiments. Data is mean ± SEM. Statistical testing was performed using a paired, two-tailed Student’s *t* test (B, E) or a two-way ANOVA (C). *, *p* < 0.05; **, *p* < 0.01; ***, *p* < 0.005; ****, *p* < 0.001; ns, not significant. Numbers in the drawn gates are percentages.

As CD226 marks CD8^+^ T cells with effector programs both early and late during TB (Fig.2, 3), we wished to further characterize how the activation state of CD226^+^ and CD226^−^ CD8^+^ T cells differ. To assess IFNγ production by T cells “in vivo” without the need for ex vivo restimulation, we infected IFNγ-IRES-eYFP (GREAT) reporter mice (*66*) with Mtb. We validated comparable expression levels of IFNγ–eYFP and IFNγ protein in response to anti-CD3 and anti-CD28 mAb stimulation in both CD4^+^ and CD8^+^ T cells, in line with published data (*67*) (Fig.S4). Although an IFNγ-reporter will miss CD8^+^ T cells expressing non-cytokine functions, we previously found that nearly all cytokine-producing CD8^+^ T cells in the lungs of Mtb-infected mice produce IFNγ (*68*). Between 10-75% of lung parenchymal CD4^+^ and CD8^+^ T cells in the lung produce IFNγ-eYFP and their frequency peaks at 23-36 wpi (Fig.4D). While both intravascular (IV^+^) and parenchymal (IV^−^) CD4^+^ T cells are IFNγ-eYFP^+^, only IV^−^ CD8^+^ T cells are IFNγ-eYFP^+^ (Fig.4D). Based on eYFP MFI, more IFNγ is made by CD226^+^ than CD226^−^ CD8^+^ T cells (Fig.4E), and when IFNγ-eYFP levels peak around 20-36 wpi (Fig.4D), higher percentages of CD226^+^CD8^+^ T cells express IFNγ-eYFP than CD226^−^CD8^+^ T cells (Fig.4E, F). Thus, using IFNγ-reporter mice, we established that CD226 identifies IFNγ-producing CD8^+^ T cells in the lungs of Mtb-infected mice in vivo.

### CD226 distinguishes phenotypically and functionally distinct CD8^+^ T cell populations

To confirm our transcriptional analysis and understand how CD226^+^ and CD226^−^ lung parenchymal CD8^+^ T cells differ functionally, we used flow cytometry to measure the expression of well-defined phenotypic markers. Compared to CD226^−^CD8^+^ T cells, CD226^+^CD8^+^ T cells express more KLRG1, TIM-3, IL-18Rα, CD103, and produce more IFNγ (Fig.5A, B, Fig.S5A). KLRG1 and CD103 expression by CD226^+^CD8^+^ T cells are mutually exclusive and, as suggested by our scRNAseq data, divides CD226^+^CD8 T cells into two differentiated subsets: KLRG1^+^ (TEFF/0) and CD103^+^ (TRM/4) (Fig.5A, B; Fig.S5B). While T-bet levels were greater in CD226^+^CD8^+^ T cells, Eomes levels were higher in CD226^−^CD8^+^ T cells, consistent with a previous finding of Eomes-dependent CD226 loss in tumors (Fig.5C, S5C) (*37*).

**Figure 5.**
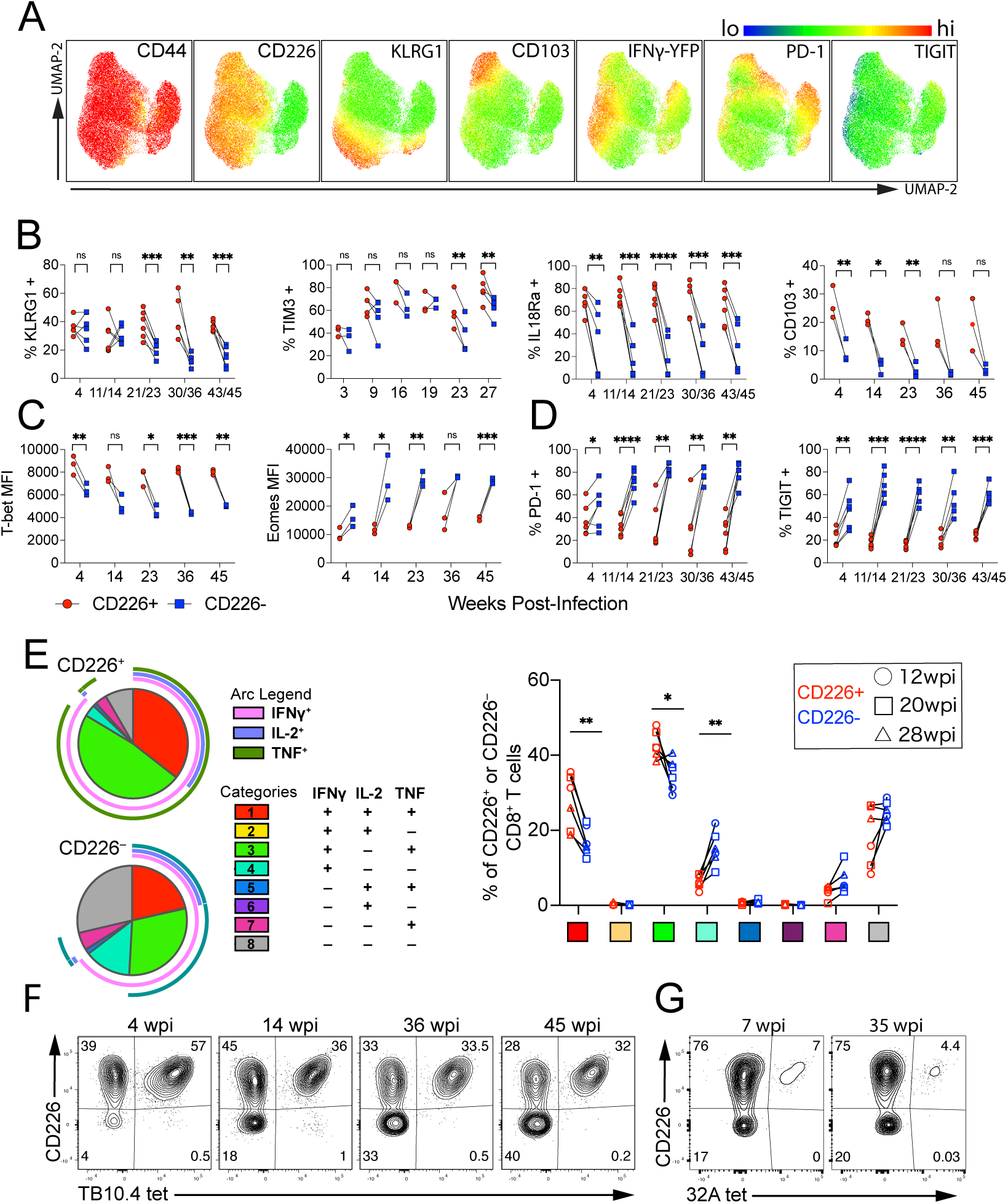
CD226^+^CD8^+^ T cells are terminally differentiated effectors with polyfunctional capacity. **(A-D)** Flow cytometric analysis of lung antigen-experienced (CD44^+^CD62L^−^) CD45-IV^−^ CD226^+^ and CD226^−^ CD8^+^ T cells from IFNγ-eYFP mice infected with Mtb. **(A)** UMAP visualization of all the antigen-experienced (CD44^+^CD62L^−^) CD45-IV^−^ CD8^+^ T cells from all combined timepoints (4, 14, 23, 36, and 45 wpi) and expression of the indicated markers were overlayed. **(B)** Quantification of the percentage of KLRG1, TIM3, IL18Rα, and CD103 expression. **(C)** Quantification of the MFI of T-bet and Eomes. **(D)** Quantification of the percentage of PD-1 and TIGIT expression. **(E)** Graph depicts the percentage of IFNγ, IL-2, and TNF expressing populations of sorted CD226^+^ and CD226^−^ lung CD45-IV^−^ CD44^+^CD8^+^ T cells stimulated with anti-CD3 and anti-CD28. Representative SPICE graph visualization at 12 wpi (left) and quantification (right). Representative flow cytometry plots of CD226 expression by TB10.4_4-11_-specific **(F)** or 32A_309-318_-specific **(G)** lung CD45-IV^−^ antigen-experienced (CD44^+^CD62L^−^) CD8^+^ T cells from IFNγ-eYFP reporter mice infected with Mtb for indicated weeks. Numbers in the drawn gates are percentages. Data from three mice per group, representative of 3 (A-D, F, G) or 2 (E) experiments. Statistical testing was performed using a paired, two-tailed Student’s *t* test. *, *p* < 0.05; **, *p* < 0.01; ***, *p* < 0.005; ****, *p* < 0.001; ns, not significant.

Over time, more CD226^−^CD8^+^ T cells express PD-1, and those that do, express higher PD-1 levels than CD226^+^ CD8^+^ T cells (Fig.5D; Fig.S5D). PD-1 expression by CD226^+^CD8^+^ T cells is limited but preferentially expressed on CD103^+^ cells (Fig.5A), consistent with enrichment of an exhaustion gene signature in TRM/4 (Fig.1F, G). When we divide CD8^+^ T cells based on PD-1 and TIM-3 expression, PD-1^−^TIM-3^+^ cells are evident after 9 wpi, and these cells have the highest CD226 levels, especially late during infection (Fig.S5E), consistent with our transcriptomic data (Fig.3C) (*29*). TIGIT expression was constantly higher in CD226^−^CD8^+^ T cells than in CD226^+^CD8^+^ T cells throughout the infection (Fig.5D, Fig.S5D).

We next compared the potential of lung parenchymal CD226^+^ and CD226^−^ CD44^+^CD8^+^ T cells to produce cytokines after ex vivo restimulation with anti-CD3 and anti-CD28 antibodies. Sorted CD226^+^CD8^+^ T cells from the lungs of Mtb-infected mice were more polyfunctional (IFNγ^+^TNF^+^IL-2^+^ or IFNγ^+^TNF^+^IL-2^−^) than CD226^−^CD8^+^ T cells, independent of the duration of infection (Fig.5E). Conversely, a higher percentage of CD226^−^CD8^+^ T cells produced only IFNγ or were IFNγ^−^TNF^−^IL-2^−^, compared to CD226^+^CD8^+^ T cells.

We next asked whether CD226 expression identifies Mtb-specific effector CD8^+^ T cells. TB10.4_4-11_ and 32A_309-318_ are immunodominant epitopes recognized by CD8^+^ T cells from C57BL/6J mice. TB10.4-specific CD8^+^ T cells maintain CD226 expression in the lung and other tissues even very late during chronic infection (Fig.5F; Fig.S5F). Similarly, nearly all 32A-specific CD8^+^ T cells express CD226 throughout the infection (Fig.5G).

Thus, cell surface CD226 expression distinguishes two major CD8^+^ T cell populations in the lung during Mtb infection. CD226^+^CD8^+^ T cells are the dominant population, which include terminally differentiated CD8^+^ T cells that express KLRG1 (TEFF/0) or CD103 (TRM/4) and are capable of polyfunctional cytokine production. In contrast, CD226^−^CD8^+^ T cells emerge as early as 4 wpi, are primarily in the TEXH/1 cluster, express PD-1, TIGIT, and Eomes, and have a reduced potential for polyfunctional cytokine production. Thus, the absence of CD226 expression by CD8^+^ T cell is an indication of CD8^+^ T cell exhaustion during TB.

### CD226 costimulates CD8^+^ T cell recognition of Mtb-infected macrophages

As CD226 identifies Mtb-specific polyfunctional CD8^+^ T cells, we sought to determine if CD226 functions in recognizing Mtb-infected macrophages. The binding of CD226 to its ligands CD155 (Poliovirus receptor, PVR) or CD112 (Nectin-2) costimulates both T cells and NK cells (*33, 69–71*). We assessed the expression of CD226 ligands by antigen-presenting cells. CD155 is expressed by nearly all alveolar macrophages (AM) and non-AM macrophages and 60% of monocytes and dendritic cells in the lungs of infected mice (Fig.6A, Fig.S6A). CD112 is expressed by 60% of alveolar and non-alveolar macrophages and 20-50% of monocytes and dendritic cells. Like lung macrophages, uninfected and infected thioglycolate-elicited peritoneal macrophages (TG-PMs), which are inflammatory recruited macrophages, expressed uniformly high levels of CD155 and lower levels of CD112 (Fig.6B).

**Figure 6.**
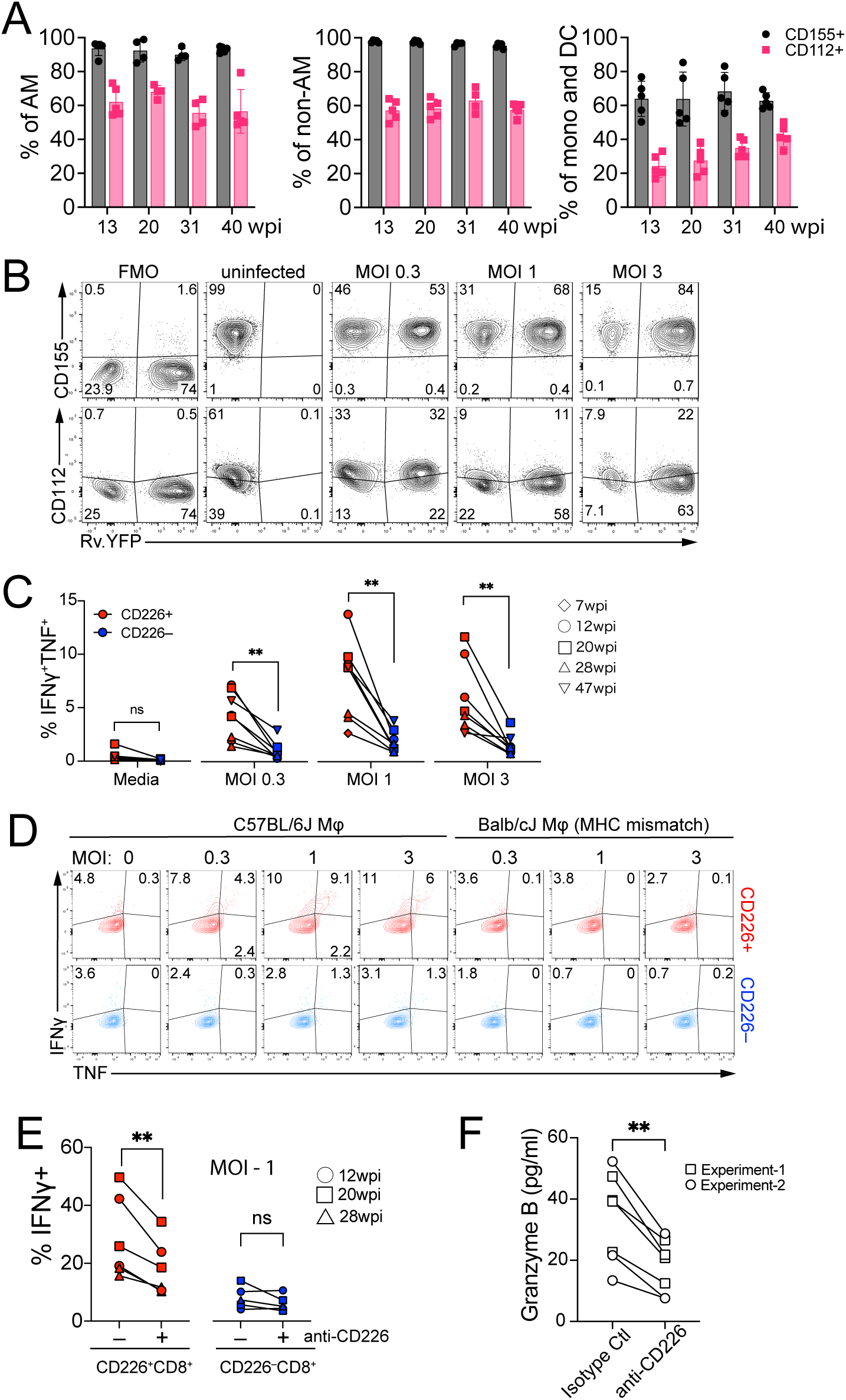
CD226 identifies and costimulates CD8^+^ T cells that recognize infected macrophages. **(A)** CD155 (grey) and CD112 (pink) expression by AM, non-AM macrophages (non-AM), monocyte and dendritic cells (mono and DC) from Mtb-infected lungs for indicated weeks. n=4-5. See Fig.S6A for gating strategy. **(B)** Flow cytometry plots showing CD155 or CD112 expression by uninfected, H37Rv-YFP infected or bystander TG-PMs. MOI, multiplicity of infection (MOI). FMO, fluorescence minus one control. (**C-E**) IFNγ and TNF production by flow-sorted lung CD45-IV^−^ CD226^+^ or CD226^−^ CD44^+^CD8^+^ T cells cultured with infected TG-PMs at the indicated MOI for 17 h. TG-PMs are from C57BL/6J (matched) or Balb/cJ (mismatched) mice. See Fig.S6B for gating strategy. Quantification (**C**) and representative flow cytometry plots of IFNγ and TNF production by CD8^+^ T cells from mice infected with Mtb for 12 weeks (**D**). **(E)** IFNγ production by flow-sorted lung CD45-IV^−^ CD226^+^ or CD226^−^ CD44^+^CD8^+^ T cells cultured with infected TG-PMs at an MOI=1 for 17 hours in the presence or absence of anti-CD226 mAb. **(F)** Granzyme B secretion by lung CD8^+^ T cells cultured with infected TG-PMs at an MOI=1 for 3 days in the presence of anti-CD226 or an isotype control mAb. Results are representative of 2-3 experiments (A, B, F) or pooled from 2 experiments (n=8) (C-E). Numbers in the drawn gates are percentages (B, D). Statistical testing was performed using a paired, two-tailed Student’s *t* test (C, E, F). *, *p* < 0.05; **, *p* < 0.01; ***, *p* < 0.001; ns, not significant.

To compare the capacity of CD226^+^ and CD226^−^ CD8^+^T cells to recognize Mtb-infected macrophages, CD45-IV^−^CD44^+^CD8^+^ T cells from the lungs of infected mice were sorted into CD226^+^ or CD226^−^ populations and cultured with infected TG-PMs (Fig.S6B). More CD226^+^ CD8^+^ T cells recognized Mtb-infected macrophages than CD226^−^ CD8^+^ T cells, based on IFNγ and TNF production measured by ICS, at all multiplicities of infection (MOI) tested (Fig.6C, D, Fig.S6C). Importantly, the recognition of Mtb-infected macrophages was specific, as there was little cytokine production when CD226^+^ CD8^+^ T cells were cocultured with MHC-mismatched Mtb-infected TG-PMs from BALB/cJ mice.

We next investigated whether CD226 is involved in the interaction between T cells and infected macrophages. Sorted lung parenchymal CD226^+^ or CD226^−^ CD44^+^CD8^+^ T cells from infected mice (12, 20, and 28 wpi) were cultured with Mtb-infected macrophages in the presence or absence of a blocking anti-CD226 mAb. Anti-CD226 mAb inhibited IFNγ production by CD226^+^ CD8^+^ T cells (Fig.6E, Fig.S6C). Additionally, CD226 blocking reduced soluble Gzmb levels detected after CD8^+^ T cell culture with infected macrophages, suggesting reduced cytotoxicity (Fig.6F). We conclude that CD226 is a marker for CD8^+^ T cells that recognize Mtb infected macrophages and functions as a costimulatory molecule in activating CD8^+^ T cells following MHC-restricted recognition of infected macrophages.

### *CD226* and *GZMK* identify distinct CD8^+^ T cell subsets from macaques and humans

We next determined if *CD226* expression is associated with effector CD8^+^ T cells in other species using published scRNA-Seq datasets of CD8^+^ T cells obtained from the lungs of Mtb-infected cynomolgus macaques and people. Previously, CD8^+^ T cells from TB granulomas in the lungs of macaques after primary infection were categorized into four subsets (*72*): 1) TEMRA-like; 2) Eff-like; 3) GZMK^hi^ TEM/PEX-like (GZMK^hi^); and 4) Tc17-like. These CD8^+^ T cells (n=3,974) were reanalyzed for *CD226* expression. *CD226* is high in TEMRA-like and Eff-like cells and lowest in GZMK^hi^ CD8^+^ T cells (Fig.7A). Reanalyzing DEGs between *CD226*^hi^ (TEMRA-like, Eff-like, and Tc17-like) and *CD226*^lo^ (GZMK^hi^) CD8^+^ T cells highlighted the mutually exclusive expression of *CD226* and *GZMK* by CD8^+^ T cells (Fig.7B, Table S6). *CD226* was associated with the expression of *CX3CR1*, which is associated with tissue residency, and the cytotoxic effector molecules *GZMA, GZMB, PRF1*, and *GNLY*; in contrast, *GZMK* was associated with *CD28, EOMES,* and *TOX* expression. Reanalysis of granuloma CD8^+^ T cells at a later time point, from another Mtb-infected cynomolgus macaque dataset found *CD226* expression to be exclusively high in cytotoxic cluster 4, which was previously identified as being associated with low-bacterial-burden granulomas (Fig.7C)(*5*). We observed the same dichotomy of *CD226* and *GZMK* expression on cytotoxic cells in reanalyzing DEGs between cytotoxic cluster 4, which has the highest *CD226* expression, and other cytotoxic clusters, which had little *CD226* expression (Fig.7D, Table S6). *CD226* was associated with the expression of *CX3CR1*, and *GZMK* was associated with the expression of *CD28*, *EOMES*, *TIGIT, GZMA,* and *GNLY* (Fig.7D). We further confirmed this dichotomic expression of *CD226* and *GZMK* on lung granuloma CD8^+^ T cells (n=1,510) from another cynomolgus dataset (*11*). We used 0.5 as a log normalized expression threshold to divide cells into CD226^hi^ (n=609) and CD226^lo^ (n=901) CD8^+^ T cells. *CD226* was associated with the expression of *IFNG* and *TNF* and the cytotoxic molecules *PRF1* and *GZMB* on CD8^+^ T cells (Fig.7E, Table S6). *GZMK* was associated with the expression of *EOMES*, *TOX*, and *TIGIT* (Fig.7E). Thus, lung granuloma *CD226*-expressing CD8^+^ T cells from infected macaques have effector features marked by the expression of *CX3CR1* and cytotoxic effector molecules and cytokines. In contrast, *GZMK* was associated with the expression of exhaustion-associated genes, including *EOMES* and *TOX*. Finally, we reanalyzed 7,241 CD8^+^ T cells from the resected lungs of patients with active TB (*73*). DEGs between CD226^hi^ (n=661) and CD226^lo^ (n=6,580) CD8^+^ T cells using 0.5 as a log normalized expression threshold to divide cells, the same dichotomy was observed between *CD226* and *GZMK* expression (Fig.7F, Table S6). *CD226*-expressing cells have enriched expression of *ZNF683*, which is related to tissue residency, and *GZMB* and *GNLY,* which encode cytotoxic effector molecules. Taken together, *CD226* and *GZMK* expression identify distinct CD8^+^ T cell subsets in the Mtb infected lung. Thus, CD226 expression identifies CD8^+^ T cells with effector features, in mice, macaques, and humans.

**Figure 7.**
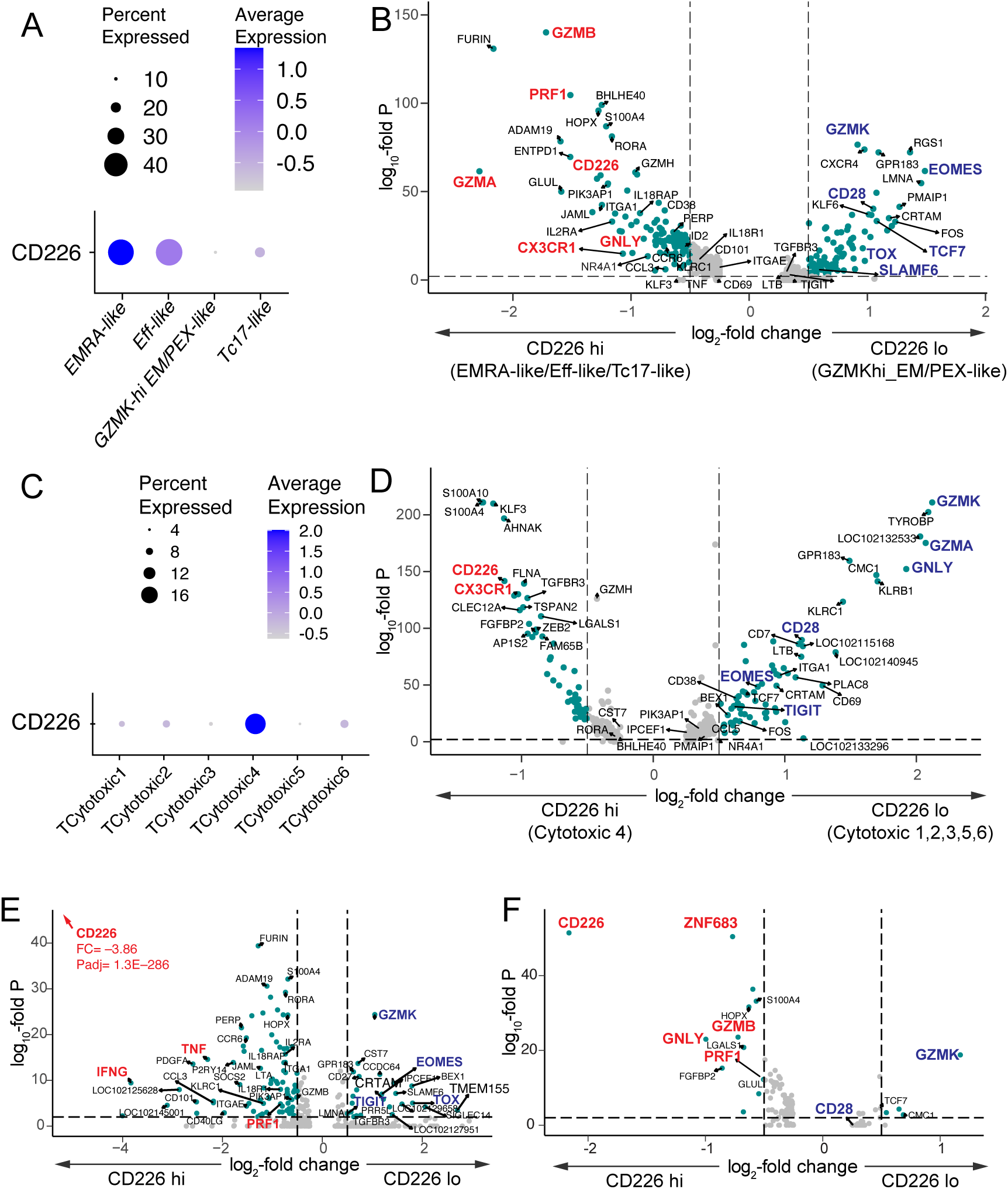
*CD226* and *GZMK* expression identify distinct lung CD8^+^ T cell subsets in Mtb infected macaque and human. **(A)** *CD226* expression by lung granuloma CD8^+^ T cell clusters during primary TB of cynomolgus macaques as defined by Bromley et al. **(B)** Volcano plots of DEGs between *CD226*^hi^ (TEMRA-like, TEFF-like, and Tc17-like) and *CD226*^lo^ (*GZMK*^hi^ TEM/PEX-like) CD8^+^ T cells during primary TB, based on reanalysis of data from Bromley et al. **(C)** *CD226* expression by lung granuloma cytotoxic cells from infected cynomolgus macaques as defined by Gideon et al. **(D)** Volcano plot of DEGs between *CD226*^hi^ (Cytotoxic 4) and *CD226*^lo^ (Cytotoxic 1,2,3,5,6) cytotoxic cells based on reanalysis of data from Gideon et al. (E) Volcano plot of DEGs between *CD226*^hi^ (log normalized expression>0.5, n=609) and *CD226*^lo^ (log normalized expression<0.5, n=901) of reanalyzed lung granuloma CD8^+^ T cells in infected cynomolgus macaques from Winchell et al. (*11*). **(F)** Volcano plot of DEGs between CD226^hi^ (log normalized expression>0.5, n=661) and CD226^lo^ (log normalized expression<0.5, n=6,580) of reanalyzed CD8^+^ T cells from resected lungs of TB patients described by Wen et al. (*73*) **(B, D, E, F)** Genes with log2FC >0.5 and an adjusted p value < 0.01 are designated by a green filled circle.

## Discussion

Recognition of Mtb-infected macrophages is crucial during CD8^+^ T cell mediated immunity during TB (*17, 23*). We previously found that some CD8^+^ T cells inefficiently recognize Mtb-infected cells by their IFNγ secretion, but the mechanisms are poorly understood (*21, 22, 24*). To determine whether heterogeneity among CD8^+^ T cells could explain the variation in recognition of Mtb-infected macrophages and inhibition of Mtb growth that is observed experimentally, we analyzed lung parenchymal (i.e., extravascular) CD8^+^ T cells by scRNA-seq, paired TCR-seq, and high dimensional flow cytometry with IFNγ-eYFP reporter mice. We identified distinct CD8^+^ T cell clusters containing clonally expanded T cells and defined functional (TEFF/0, TRM/4, and TPOLY/6) and dysfunctional (TEXH/1 and TIFN/8) states, at 6- and 41-weeks after infection. An important discovery is that CD226 expression distinguishes effector CD8^+^ T cells with retained polyfunctionality from dysfunctional CD8^+^ T cells throughout Mtb infection. Importantly, CD8^+^ T cell with dysfunctional states, characterized by the lack of CD226 expression, did not recognize infected macrophages. On the other hand, CD8^+^ T cells with functional states characterized by CD226 expression, recognized infected macrophages in an MHC-restricted manner, and blocking CD226 diminished recognition. CD226 is a T cell activation marker, functions as an adhesion molecule, and costimulates T cells (*33, 69, 70, 74*). Costimulation occurs when CD226 binds to CD155 or CD112 (*71*). Our work highlights a novel role of the CD226 costimulatory signal in enhancing CD8^+^ T cell recognition of Mtb-infected macrophages.

CD226 signaling by tumor-infiltrating CD8^+^ T cells is negatively regulated by PD-1 and TIGIT(*75*). PD-1 inhibits the phosphorylation of CD226 via its ITIM-containing intracellular domain (*75*). TIGIT competes with CD226 for binding to CD155, and ligation to CD155 induces internalization and degradation of CD226 as a possible negative feedback mechanism (*35*). The loss of the expression or signaling of CD226 restrains CD8^+^ T cell functions, leads to the accumulation of dysfunctional CD8^+^ T cells in the tumor microenvironment, and is associated with impaired tumor control (*35*). In the setting of cancer, PD-1 and TIGIT are often co-expressed with CD226 on CD8^+^ T cells, and CD226 expression is required for response to anti-PD-(L)1 or anti-TIGIT antibody treatment (*36, 37, 75*). However, during chronic TB, PD-1 and TIGIT are expressed at low levels on CD226^+^ CD8^+^ T cells. Dysfunctional CD8^+^ T cells that have entirely lost CD226 expression now express high levels of PD-1 and TIGIT. This distinct expression pattern of PD-1, TIGIT, and CD226 might explain distinct outcome induced by PD-1 blockade in TB (*76*). In a murine cancer model, additional mechanisms occur at the transcriptional level to restrict CD226 expression, where Eomes promote CD226 loss by tumor-infiltrating CD8^+^ T cells (*37*). In our study, like in cancer, CD226^+^CD8^+^ T cells have high T-bet and low Eomes expression, while CD226^−^CD8^+^ T cells have low T-bet and high Eomes levels, suggesting that balance of T-bet and Eomes affects CD226 expression during TB (*37*).

Surprisingly, we found a distinct TIM-3 expression pattern on CD8^+^ T cells during Mtb infection. While TIM-3^+^PD-1^−^ CD8^+^ T cells are not detected in cancer or chronic LCMV infection, in chronic Mtb infection, nearly 25% of the CD8^+^ T cells are TIM-3^+^PD-1^−^, and this subset has high CD226 levels and an effector state. This is consistent with TIM-3 signaling driving stronger effector functions in CD4^+^ and CD8^+^ T cells in Mtb infected mice(*29*) and active TB patients (*77*). TIM-3 is recruited to the immune synapse and enhances phosphorylation of ribosomal S6 protein and Akt/mTOR signaling under acute stimulation such as during *Listeria monocytogenes* infection (*78–81*). Sustained expression of TIM-3 with PD-1 is associated with terminally exhausted CD8^+^ T cells during cancer and chronic LCMV infection (*65, 82*). While robust TIM-3 upregulation in the absence of PD-1 on CD8^+^ T cells is also observed during *Mycobacterium avium* infection (*83*), and TIM-3^+^PD-1^−^ CD8^+^ T cells are also found in HIV-1 and HCV infection (*84, 85*), the functionality of TIM-3^+^PD-1^−^ CD8^+^ T cells are poorly described. Our data support the costimulatory role of TIM-3 to help effector function in the absence of PD-1 during Mtb infection.

There are parallels between how CD226 and CD28 costimulate T cells and are regulated. Both are immunoglobulin superfamily members that bind ligands on APCs (CD155 or CD112, and CD80 and CD86, respectively). Coinhibitory receptors compete for ligand binding as a mechanism to down-regulate or terminate T cell responses. TIGIT competes for CD155 binding, while CTLA4 competes for CD28 binding. Both CD226 and CD28 are regulated by PD-1 signaling (*75*). These mechanistic insights led to the discovery that successful checkpoint blockade targeting PD-1 and TIGIT during cancer depends on the presence of CD226^+^CD8^+^ T cells (*36, 37, 75*). CD226 and CD28 have opposite expression patterns in two cynomolgus macaque datasets. Thus, CD226 could also be an important costimulatory molecule for macaque CD8^+^ T cells during Mtb infection. Finally, we find an interesting dichotomy between CD226 and GZMK expression among CD8^+^ T cells. Although GZMK, in combination with perforin, has cytolytic and anti-mycobacterial activity in vitro (*86*), in our dataset, it is associated with the two dysfunctional clusters, TEXH/1 and TIFN/8. In scRNA-Seq data from all species (mouse, cynomolgus macaque and human), *GZMK* is expressed by CD226^lo^CD8^+^ T cells. In Mtb-infected cynomolgus macaques, a population of CD226^lo^CD8^+^ T cells express high *GZMK* levels together with *TIGIT, EOMES, TOX,* and *CD28* (*72*). Interestingly, a higher proportion of lung granuloma cytotoxic 4 cells from Gideon *et al.* (*5*), which have exclusive expression of *CD226*, were associated with Low-bacterial-burden granulomas, but cytotoxic 5 cells, which are distinguished by elevated expression of *GZMK*, did not associate with bacterial control (*5*). The functional role of GZMK-expressing CD8^+^ T cells during TB requires further investigation.

We do not yet know whether CD226 expression on CD8^+^ T cells promotes protection against TB. CD226 blockade in vitro diminishes IFNγ and granzyme B production by CD8^+^ T cells. If CD226 costimulates cytokine production and cytotoxic functions of CD8^+^ T cells in vivo, we predict that CD226 signaling would reduce the survival of intracellular Mtb in macrophages. As CD226 is involved in priming (*70*), it might have a role in the robust expansion and maintenance of CD226^+^CD8^+^ T cells during infection. Further study using CD226 knockout mice to investigate its role in priming, expansion, and control against infected cells can elucidate its contribution to protective CD8^+^ T cell immunity in TB. In conclusion, we identified CD226 as a functional marker that identifies effector CD8^+^ T cells with the capacity to recognize infected macrophages. We predict that CD226 will identify effector CD8^+^ T cells in cynomolgus macaques and humans based on analysis of transcriptional data. As CD226 is lost by dysfunctional CD8^+^ T cells, changes in its expression might be useful in evaluating impaired CD8^+^ T cell responses and disease progression in clinical settings. Finally, the CD226-CD155/CD112 axis should be considered as a strategy to enhance immune responses induced by vaccination (*87*).

## Methods

### Ethics statement

Studies were conducted using the relevant guidelines and regulations and approved by the Institutional Animal Care and Use Committee at the University of Massachusetts Medical School (UMMS) (Animal Welfare A3306-01), using the recommendations from the Guide for the Care and Use of Laboratory Animals of the National Institutes of Health and the Office of Laboratory Animal Welfare.

### Mice

C57BL/6J, BALB/cJ and IFNγ-IRES-eYFP reporter (GREAT) mice (*66*) (Strain #017580) were purchased from The Jackson Laboratory. The mice used in this study were 6–8-week-old in the animal facility at the UMass Chan Medical School. Mice of both sexes were used and the experiments reported herein did not test sex as a variable.

### Mtb strain

The Erdman strain was used for aerosol infection (*88*). H37Rv-pLux or Rv.YFP strain (H37Rv expressing yellow fluorescent protein (YFP)(*89*) were used to infect macrophages *in vitro*.

### Aerosolized Mtb infection of mice

Mice were infected by the aerosol route as previously described(*89*). Briefly, frozen bacterial stocks were thawed and added to 3 ml of 0.01% Tween-80 in PBS. The bacteria were sonicated for 1 minute, and 2 ml of 0.01% Tween-80 in PBS were added. To infect mice, the bacterial suspension was aerosolized using a Glas-Col chamber (Terre Haute). The average number of bacteria delivered into the lung was determined for each experiment by plating lung homogenate on 7H11 plates (Hardy Diagnostics) from 5 mice within 24 hours after infection and ranged between 50-200 CFU/mouse.

### Cell isolation

Mtb-infected C57BL/6J mice were injected intravenously with 2.5 μg of fluorochrome-labeled anti-CD45.2 3 minutes before euthanasia and lung removal. Single cell suspensions were prepared by homogenizing lungs using a GentleMACS tissue dissociator (Miltenyi), digesting with 300 U/ml Type IV Collagenase (Sigma-Aldrich, C5138-5G) in complete RPMI (RPMI-1640 medium supplemented with 10% FBS, 1× Non-Essential Amino Acids (Gibco, 11140050) and 10 mM HEPES (Gibco, 15630080), 2 mM L-Glutamine (Gibco, 25030081), 100 units/ml Penicillin– Streptomycin (Gibco, 15140122), 0.5X MEM Amino Acids (Gibco, 11130051), and 55μM 2-Mercaptoethanol (Gibco, 21985023)) at 37°C for 30 minutes, and followed by a second run of dissociation using the GentleMACS. Suspensions were filtered through 70-μm strainers, and red blood cells were lysed in ACK Lysis Buffer (Gibco; Thermo Fisher Scientific). Suspensions were then filtered through 40-μm strainers. Mediastinal LN or spleen were harvested, mashed with a 3-ml syringe on a 24-well plate with RPMI 1640, filtered with 70-μm strainers, washed with RPMI-1640, and resuspended with autoMACS running buffer (Miltenyi).

### Flow cytometry

Single-cell suspensions, prepared as described earlier in Cell isolation section, were stained with Zombie Fixable Viability dye (Biolegend) in PBS for 10 minutes at room temperature (RT). After washing cells with autoMACS running buffer, cells were incubated with anti-mouse CD16/32 (BioXcell) for 5 minutes, followed by surface staining performed for 20 min at 4°C in autoMACS running buffer. TB10.4_4-11_ and/or 32A_309-318_ tetramers were stained together with antibodies for surface staining. The following antibodies were used: Anti-mouse CD90.2 (53-2.1), CD8b (YTS156.7.7), CD8a (53-6.7), CD4 (RM4-5 or GK1.5), CD45R/B220 (RA3-6B2), CD3 (17A2), CD44 (IM7), TIGIT/Vstm3 (1G9), CD279/PD-1 (29F.1A12), CD366/Tim-3 (RMT3-23), NK-1.1 (PK136), CD155/PVR (TX56), CD226 (10E5), CD218a/IL-18Rα (A17071D), TCRβ chain (H57-597), KLRG1/MAFA (2F1/KLRG1), CD103 (2E7), CD11c (N418), CD11b (M1/70), MERTK/Mer (2B10C42), Ly-6G (1A8), CD64 (X54-5/7.1), CD19 (1D3/CD19), IFN-γ (XMG1.2), TNF-α (MP6-XT22), IL-2 (JES6-5H4), T-bet (4B10) (all from BioLegend). Anti-mouse Siglec-F (E50–2440), CD45.2 (104), CD112/Nectin-2 (829038), CD62L (MEL-14), γδ T-Cell Receptor (GL3) were obtained from BD Biosciences, and Eomes (Dan11mag) was obtained from Invitrogen. After washing with autoMACS running buffer, stained cells were fixed in 1% paraformaldehyde. Samples were assessed on a Cytek Aurora (Cytek). Flow cytometry data were processed and analyzed using FlowJo software (BD bioscience) and FlowJo plugins UMAP v4.1.1. Lung parenchymal CD8^+^ T cells were defined as intravascular (IV) CD45^−^, CD90.2^+^CD4^−^CD8α^+^CD8β^+^. 2,294 lung parenchymal antigen-experienced (CD44^+^CD62L^−^) CD8^+^ T cells from each mouse (n=3/time point) were used for concatenation to one FCS file before creating UMAP projections.

### Cell Sorting

Cells were prepared as described earlier in Cell isolation section. Dead cells were removed using EasySep™ Dead Cell Removal (Annexin V) Kit (Stemcell) before staining. Cell staining was performed as described earlier in the Flow cytometry section. Stained cells were sorted by MA900 Multi-Application Cell Sorter (Sony).

### scRNA-seq library preparation

Lung single cell suspensions were prepared as above in Cell isolation section, and T cells from individual C57BL/6J mice were sorted based on CD45-IV^−^CD3^+^TCRβ^+^ (n=2 at 6 wpi; n=2 at 41 wpi). Antibodies for NK1.1, CD19 and TCRγδ were used for the “dump” channel. 10,000 flow-sorted T cells from mouse were loaded into a single Chip channel (10× Chromium). Gel beads-in-emulsion (GEM) generation, cDNA amplification, and library construction were performed using Chromium Single Cell 5’ Library Construction Kit (v2 Chemistry Dual Index). Chromium Single Cell V(D)J Enrichment Kit, Mouse T Cell were used to generate TCR Libraries. The final libraries were sequenced on the Illumina NextSeq 500 platform.

### scRNA-seq data processing and analysis

Sequencing reads were mapped to mouse reference genome GRCm38 (mm10) and processed through CellRanger (v6, 10× Genomics). All downstream analyses were performed through Seurat (v.4.1.1) for R (v4.2.0). Low-quality cells where >10% of transcripts derived from the mitochondria were excluded. The data was filtered to retain cells with 300-5,000 genes detected and UMI counts of 500-25,000. The TCR genes (Tra[vjc], Trb[vdjc], Trg[vj], Trd[vdjc]) were removed for from the gene expression dataset to avoid any TCR gene-driven bias during clustering. CD8^+^ T cells were selected based on Cd4 < 1e-10 AND Cd8a > 0.5 AND Cd8b1 > 0.5 AND Cd3e > 1. SCTransform normalization was performed. Linear dimensional reduction was performed using *RunPCA* with argument npcs = 30, *RunUMAP*, *FindNeighbors,* and *FindClusters* with argument resolution = 0.9. DoubletFinder (v2.0.3), assuming a theoretical doublet rate of 0.8-4.6% depend on the samples, was additionally used to remove doublets. Resulting doublets were 0.3-4.1%. To combine data across all samples, highly variable 3000 genes were selected for anchoring using the *SelectIntegrationFeatures* function, and integration was performed using *PrepSCTIntegration*, *FindIntegrationAnchors,* and *IntegrateData* functions. *RunUMAP* was performed on the integrated dataset, followed by *FindNeighbors* (reduction = “pca”) using the first 10 principal components. *FindClusters* (resolution = 0.3) was performed, and 9 clusters were identified. The cluster markers were found using the *FindAllMarkers* function (min.pct = 0.25, assay = “RNA”), and avg_log2 fold change (FC) and adjusted P value are shown in Table S1. Barplot was visualized by dittoseq (v1.10.0). Volcanoplots were created with the R package EnhancedVolcano (v1.16.0).

### Gene signature scoring

We computed average gene expression scores for published gene sets with *AddModuleScore* function in Seurat (v.4.1.1).

### Projection of published scRNA-seq data to our scRNA-seq data

Published processed Seurat object (*58*) was used as a reference to apply label transfer to our dataset using *ProjecTILs.classifier* function in ProjecTILs (v3.3.0).

### Pseudotime trajectory analysis

To determine the potential development lineages of CD8^+^ T cell clusters, we used Monocle3 (v.1.3.1) on the integrated Seurat object. Using the SeuratWrappers package, we converted the integrated Seurat object to a cell dataset object. We selected the naive T cell cluster (TN/3) as the root for the trajectory. Cells were clustered with *cluster_cells* using “Louvain” as a cluster_method. Two separate partitions for the clusters were detected, and the partition with the proliferating CD8^+^ T cells (T_PROLIF_/7) was excluded from the pseudotime trajectory analysis. Trajectory graph learning and pseudotime measurement were performed using *learn_graph* and *order_cells* function. Next, we used the *graph_test* function to identify genes that are differentially expressed on different paths through the trajectory. The genes that had a significant q-value (<0.05) from the autocorrelation analysis were grouped into 49 distinct co-regulated modules using *find_gene_modules* function (resolution=1e-2) (Table S5). The *aggregate_gene_expression* function was used to calculate aggregate expression of genes in each module for all the clusters. We used the *enrichGO* function in the clusterProfiler package (v 4.6.2) to measure the enrichment of the modules in GO terms across all three ontologies (“BP,” “MF,” and “CC”).

### Cytokine signaling activity score

The cytokine signaling activity of each Seurat cluster on the integrated scRNA-Seq data was predicted using CytoSig(*53*). Averaged gene expression on 2000 variable genes of all cells in each Seurat cluster was calculated using *AverageExpression* function (assays = “RNA”). Predicted cytokine signaling activity scores were calculated in CytoSig (https://cytosig.ccr.cancer.gov/) and plotted in a heatmap using the R package ComplexHeatmap (v2.14.0)

### SCENIC regulons

R package SCENIC (v1.3.1) was used for GRN inference. For efficient analysis, the integrated scRNA-Seq dataset was down-sampled to 3000 cells. We created the initialize settings configuration object with *initializeScenic* function with default settings. To remove genes that are expressed either at very low levels or in too few cells, we only kept the genes with at least 6 UMI counts, detected in at least 1% of the cells, and are available in RcisTarget databases. After calculating the spearman correlation with *runCorrelation* function, we used *runGenie3* function to infer potential transcription factor targets. For building the gene regulatory network, transcriptional factors and their top 10 potential gene targets were predicted with the following SCENIC functions: *runSCENIC_1_coexNetwork2modules*, *runSCENIC_2_createRegulons,*and *runSCENIC_3_scoreCells*. Regulon activity for each cell was calculated as the average normalized expression of putative target genes. Regulon Activity were plotted in a heatmap using the R package ComplexHeatmap (v2.14.0). All required inputs were downloaded from https://resources.aertslab.org/cistarget/databases/old/mus_musculus/mm9/refseq_r45/mc9nr/gene_based/.

### Cynomolgus macaque scRNA-seq analysis

For the cynomolgus macaque granuloma dataset from Bromley *et al.* (*72*), we downloaded the processed scRNA-seq object from the Broad Single Cell Portal. The dataset was filtered to include only CD8^+^ T cells which are “TEMRA-like,” “GZMK^hi^ TEM/PEX-like,” “TEff-like,” and “Tc17-like” on SubclusteringV2. Primary infection group was selected based on Naïve in the Group_Detailed category. For the cynomolgus macaque granuloma dataset at 10wpi from Gideon *et al.* (*5*), we downloaded raw scRNA-seq data from GSE200151. The dataset was filtered to include only cytotoxic cells (“TCytotoxic1,” “TCytotoxic2,” “TCytotoxic3,” “TCytotoxic4,” “TCytotoxic5,” and “TCytotoxic6”) on SpecificFinal category. *NormalizeData*, *ScaleData*, *FindVariableFeatures*, and *RunPCA* were performed by Seurat (v.4.1.1) with default settings. For the cynomolgus macaque granuloma dataset from Winchell *et al.* (*11*), we downloaded processed scRNA-seq data from the Broad Single Cell Portal. The dataset was filtered to include T/NK cells in the General_Celltypes category, and then CD8^+^ T cells were further selected based on CD8A > 0.5 AND CD8B > 0.5. Only IgG treated group was used for further analysis. We used 0.5 as a log normalized expression threshold to divide cells into CD226^hi^ (n=609) and CD226^lo^ (n=901) CD8^+^ T cells. For all datasets, differentially expressed gene analysis was performed between indicated CD8^+^ T cell populations using the *FindMarkers* function and visualized using EnhancedVolcano (v1.16.0) and ggplot2 (v3.5.1). DEGs are listed in Table S6.

### Human scRNA-seq analysis

scRNA-seq data from resected lungs from active TB patients were downloaded from Wen *et al.* and QC and processed as described earlier in scRNA-seq data processing and analysis. We excluded a sample of SP020L due to the small cell number when integration was performed. The dataset was filtered to only include CD8^+^ T cells based on CD8A > 0.5 AND CD8B > 0.5 AND CD3E > 1 AND CD4 < 1e-10. We used 0.5 as a log normalized expression threshold to divide cells into CD226^hi^ (n=661) and CD226^lo^ (n=6,580) CD8^+^ T cells. Differentially expressed gene analysis was performed between CD226^hi^ and CD226^lo^ CD8^+^ T cells using the *FindMarkers* function and visualized using EnhancedVolcano (v1.16.0) and ggplot2 (v3.5.1). DEGs are listed in Table S6.

### CD8^+^ T cell clonotype filtering and analysis

Cell Ranger VDJ pipeline (v6, 10× Genomics) was used to process the raw TCR sequence data with default augments and align them to the mouse reference genome GRCm38 (mm10). CD8^+^ T cells were filtered based on *Cd3e* (>1) AND *Cd8a* AND *Cd8b1* (>0.5) AND *Cd4*< 1e-10 with a single rearranged TCR comprised of a single TCRα and TCRβ chain. CD8^+^ T cells that met these criteria were then assigned to a clonotype using the nucleotide sequences of the CDR3 region of TCR α and β. Repertoire overlap analysis was performed using R package Immunarch (v1.0.0), using *repOverlap* function with .method = “morisita” and .col = “nt”. Because of the large imbalance of cell number in each Seurat cluster, we uniformly down-sampled all clusters to 209 cells for clonal overlap calculation. TRV and TRJ repertoires and CDR3 sequences for all TCR clonotypes are listed in Table S2.

### RNA Sequencing and data analysis

CD226^+^ or CD226^−^ CD45-IV^−^CD44^+^CD8^+^ T cells were isolated by flow sorting from 3 C57BL/6J mice each, which were infected with Mtb for 24-28 weeks prior to the sort. Total RNA was extracted from sorted cells stored in RNAprotect (QIAGEN) using Direct-zol RNA Microprep kit (Zymo research) according to the manufacturer’s instructions. RNA sequencing with poly(A) selection was performed at GENEWIZ/AZENTA. Sequence reads were trimmed to remove possible adapter sequences and nucleotides with poor quality using fastp v.0.23.1. UMI-based de-duplication was performed using fastp (v.0.23.1) simultaneously. Trimmed and de-duplicated reads were then mapped to the Mus musculus GRCm38 reference genome available on ENSEMBL using the STAR aligner (v.2.5.2b). Normalization and differential gene expression analysis were performed using DESeq2 R package (v1.38.3). The raw read matrix was log2 transformed using the *rlogTransformation* function, and Principal component analysis (PCA) was created. For GSEA, we used fgsea (v1.18.0) with MSigDB (v7.5.1) immunologic.signature_gene_sets (C7).

### Published RNA-Sequencing and Nanostring data analysis

Normalized gene expression values from published RNA-seq data (*63, 64*) and Nanostring data (*29*) were obtained, and Z-scaled scores were shown in a heatmap using the R package pheatmap (v1.0.12).

### Intracellular cytokine staining

Cells were incubated in complete RPMI at 37°C/5% CO2 for 17 hours in the presence or absence of stimulation. In some experiments cells were stimulated with anti-CD3 (1 μg/ml) and anti-CD28 (5 μg/ml) for 5 hours. During the last 5 hours of incubation, we added Brefeldin A (eBioscience) at final concentration of 3 ug/ml. Cells were stained with Zombie Fixable Viability dye (Biolegend) and surface staining was performed as described above. Cells were then fixed and permeabilized for 20 minutes at 4°C using BD Cytofix/Cytoperm permeabilization kit (BD Biosciences). Cells were then washed with Perm/Wash buffer and stained with the intracellular antibodies in Perm/Wash buffer for 30 minutes at 4°C. After washing with autoMACS running buffer, stained cells were fixed in 1% paraformaldehyde. Data was collected on a Cytek Aurora (Cytek). Polyfunctionality pie charts were created using SPICE 6.1 provided by the National Institute of Allergy and Infectious Diseases (NIAID).

### Transcription factor staining

Cells were stained with Zombie Fixable Viability dye (Biolegend) and surface staining was performed as described above. Cells were then fixed and permeabilized with the FoxP3/transcription factor buffer staining set (eBioscience) for 20 minutes at RT. Cells were washed with 1X Permeabilization Buffer and stained with antibody cocktail diluted in 1X Permeabilization Buffer for 30 minutes at RT. After washing with autoMACS running buffer, stained cells were fixed in 1% paraformaldehyde. Samples were assessed on a Cytek Aurora (Cytek).

### Preparation of Mtb

Mtb was grown in 7H9 media (supplemented with 10% OADC, 0.05% of Tween-80, and 0.2% glycerol) until an OD_600_ = 0.6-0.8. The bacteria were washed with RPMI 1640, and incubated with TB coat (RPMI 1640 containing 1% heat-inactivated FBS, 2% human serum, and 0.05% Tween-80) at RT for 5 minutes. After washing again with RPMI 1640, the bacteria passed through a 5 μm filter to remove clumps. OD_600_ was measured again to adjust the concentration with a conversion factor of OD_600_ 1 = 3 · 10^8^ bacteria/ml, providing a multiplicity of infection (MOI) of 0.3, 1, 3 in cRPMI (without antibiotics).

### Collecting of thioglycolate-elicited peritoneal macrophages (TG-PMs) and in vitro infection

Thioglycolate was injected into the peritoneal cavity in C57BL/6J or BALB/c mice. After 4 days, peritoneal lavage was collected from peritoneal cavity, and macrophages were purified using CD11b microbeads (Miltenyi). Purified TG-PMs were plated 10^5^/well in 96 well flat plates and, once adhered, infected with Mtb overnight at 37°C/5% CO2. TG-PMs infected at MOI 1 were lysed with 1% Triton X-100 the next day (Day 1) and plated with serial dilutions of the lysate on 7H11 plates (Hardy Diagnosis). The plates were incubated for 21 days at 37°C/5% CO2 for Day 1 CFU enumeration.

### In vitro coculture of sorted CD8^+^ T cells and infected TG-PMs

TG-PMs were infected in vitro overnight or left uninfected as described above. The next day, the TG-PMs were washed three times with RPMI1640 to remove any extracellular bacteria. Sorted CD226^+^ or CD226^−^ CD45-IV^−^CD44^+^CD8^+^ T cells were added to the TG-PMs at a ratio of 1:2 (T cell to macrophage). The cells were cultured in cRPMI (without antibiotics) at 37°C/5% CO2 for 17 hours in the presence or absence of 25 μg/ml anti-CD226 mAb (10E5, eBioscience). Intracellular cytokine staining was performed as described above.

### Cytokine measurements in coculture supernatant

Single cell suspensions were prepared from the lungs of Mtb-infected C57BL/6J mice as described above using GentleMACS tissue dissociators (Miltenyi). Cell suspensions were filtered through 70-μm strainers, and red blood cells were lysed in ACK Lysis Buffer (Gibco; Thermo Fisher Scientific). Cell suspensions were then filtered through 40-μm strainers. CD8^+^ T cells were purified from suspensions using mouse CD8 (TIL) MicroBeads (Miltenyi), resulting in highly pure products (>92% CD8^+^). Purified CD8^+^ T cells were cocultured with Mtb-infected TG-PMs (MOI=1) in the presence of anti-CD226 (10E5) or isotype control mAb, both at 25 μg/ml, for 72 hours in cRPMI (without antibiotics) at 37°C/5% CO2. Cell culture supernatant was collected, and mouse Granzyme B was quantified via ELISA (Mouse Granzyme B DuoSet ELISA, R&D Systems) following manufacturer’s instructions.

### Statistical analysis

Statistical analyses were performed using GraphPad Prism (v10) software. The statistical tests performed for each experiment are specified in the figure legends.

### Data availability

Sequencing data generated for this study have been deposited in the Gene Expression Omnibus database with accession code GSE266006.

## Acknowledgements

We thank the UMass Chan Flow Cytometry Core for their expertise and NIH S10OD028576 for the purchase of the BD FACSFusion Cell Sorter. Tetramers were produced by the NIAID Tetramer Core (Emory, Atlanta, Georgia).

## Author contributions

Conceptualization: TS, SMB

Investigation: TS, EC, KC, RL, TR

Formal analysis: TS, RL, SMB

Writing & Editing: TS, SMB, RL, TR

Supervision: SMB

Funding Acquisition: SMB

## Data and materials availability

Sequencing data generated for this study have been deposited in the Gene Expression Omnibus database with accession code GSE266006. All other data are available in the main text or the supplementary materials.

## Supplementary Materials

**Supplemental Figure 1.** Nine distinct lung parenchymal CD8^+^ T cell transcriptional states are identified during Mtb infection.

**Supplemental Figure 2.** Transcriptional analysis of CD226^+^CD8^+^ T cells demonstrates expression of an effector program.

**Supplemental Figure 3.** CD226 expression on lung immune cell subsets.

**Supplemental Figure 4.** IFNγ-eYFP expression correlates with intracellular IFNγ measured by ICS on T cells from IFNγ-eYFP mice.

**Supplemental Figure 5.** CD226 expression identifies terminally differentiated effector CD8^+^ T cells.

**Supplemental Figure 6.** Representative gating strategies for myeloid cells and T cells.

**Table S1.** DEGs expressed by each cluster in scRNA-seq of lung parenchymal CD8^+^ T cells from 6 and 41wpi. 3 tabs corresponds top 30 upregulated genes (tab1), DEGs with average log2 fold change >0.5 or <-0.5, and adjusted P value < 0.01 (tab2) and DEGs with average log2 fold change >0.1 or <-0.1 and adjusted P value < 0.01 (tab3) in each cluster.

**Table S2.** scTCR-Seq of lung parenchymal CD8^+^ T cells from 6 and 41wpi.

**Table S3.** DEGs between TEFF/RM branch [TEFF/0, TSL-M/2, TRM/4, and TPOLY/6] and TEXH branch [TEXH/1 and TIFN/8] (log2FC > 0.5 or <-0.5, adjusted P value ≤ 0.01), related to Figure 2D.

**Table S4.** DEGs between lung parenchymal (CD45-IV^−^) CD226^+^ and CD226^−^ CD44^+^ CD8^+^ T cells from mice with chronic infection (24-28wpi) analyzed by RNA-Seq, related to Figure 3A, B and Figure S2A.

**Table S5.** List of modules of coregulated genes identified by Monocle3 in scRNA-seq of lung parenchymal CD8^+^ T cells from 6 and 41wpi, related to Figure S2B, C. The genes that had a significant q-value (<0.05) from the autocorrelation analysis were grouped into 49 distinct co-regulated modules.

**Table S6**. DEGs between CD226^hi^ and CD226^lo^ CD8^+^ T cells in reanalyzed published datasets, related to Figure 7.

