## Supplemental Figures for "CD226 identifies effector CD8^+^ T cells during tuberculosis and costimulates recognition of *Mycobacterium tuberculosis*-infected macrophages"

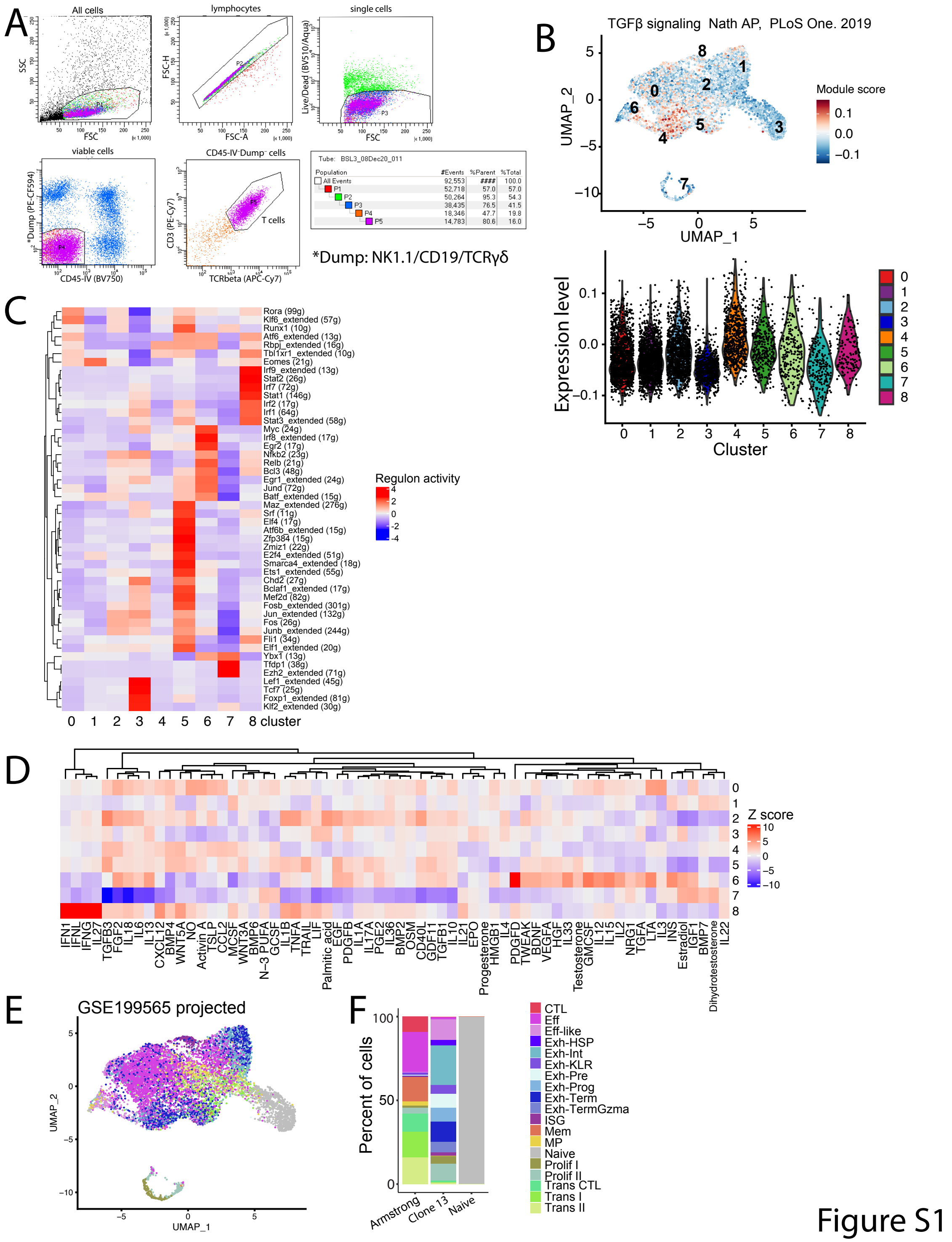


**Supplemental Figure 1. Nine distinct lung parenchymal CD8$^{+}$ T cell transcriptional states are identified during Mtb infection.** **(A)** Gating strategy for flow cytometric sorting of lung parenchymal CD45-IV$^{-}$ CD3$^{+}$ TCRβ$^{+}$ T cells for scRNA-seq and paired scTCR-seq. Antibodies for NK1.1, CD19, and TCRγδ were used for the "dump" channel. **(B)** UMAP (upper panel) and violin plot (bottom panel) depicting the enrichment of TGFβ signature(*42*) (log2FC>0.5; adjusted P value < 0.05). **(C)** Heatmap showing per cell scaled regulon activity scores (presented as z-score)

as inferred by SCENIC algorithm for each cluster. **(D)** Heatmap showing activity scores (presented as z-score) of the response to different cytokines and chemokines quantified by Cytosig for each cluster**. (E)** UMAP showing projection of gp33-specific $CD8^+$ T cells from spleens of mice infected with LCMV [GSE199565](*58*) onto our $CD8^+$ T cell dataset from 6 and 41wpi. **(F)** Reanalyzed stacked barplots showing the proportions of phenotypic clusters among splenic $CD8^+$ T cells from Armstrong-infected, Clone 13-infected or uninfected mice [GSE199565](*58*).

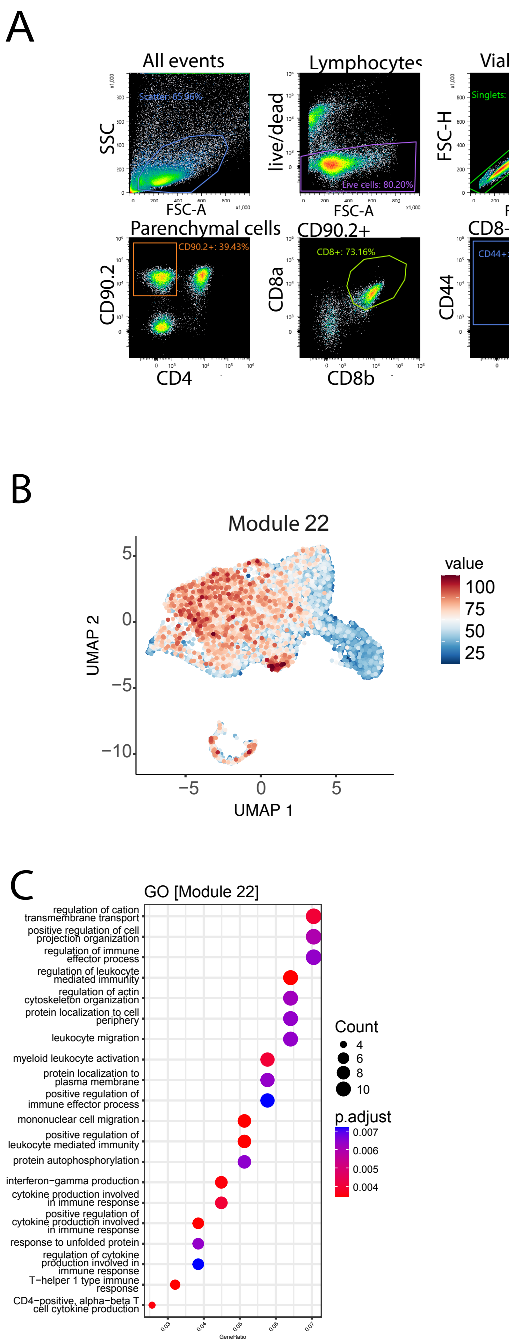


**Supplemental Figure 2. Transcriptional analysis of CD226$^+$CD8$^+$ T cells demonstrates expression of an effector program. (A)** Lung parenchymal CD45-IV$^-$ CD226$^+$ and CD226$^-$ CD44$^+$ CD8$^+$ T cells from mice infected with Mtb for 24-28 weeks were analyzed by RNA-seq. 3 mice were pooled for each triplicate sample. Gating strategy for flow cytometric sorting analysis. **(B)** UMAP showing module 22 gene expression from Monocle3's co-regulated gene module analysis. **(C)** Gene ontology term analysis of module 22.

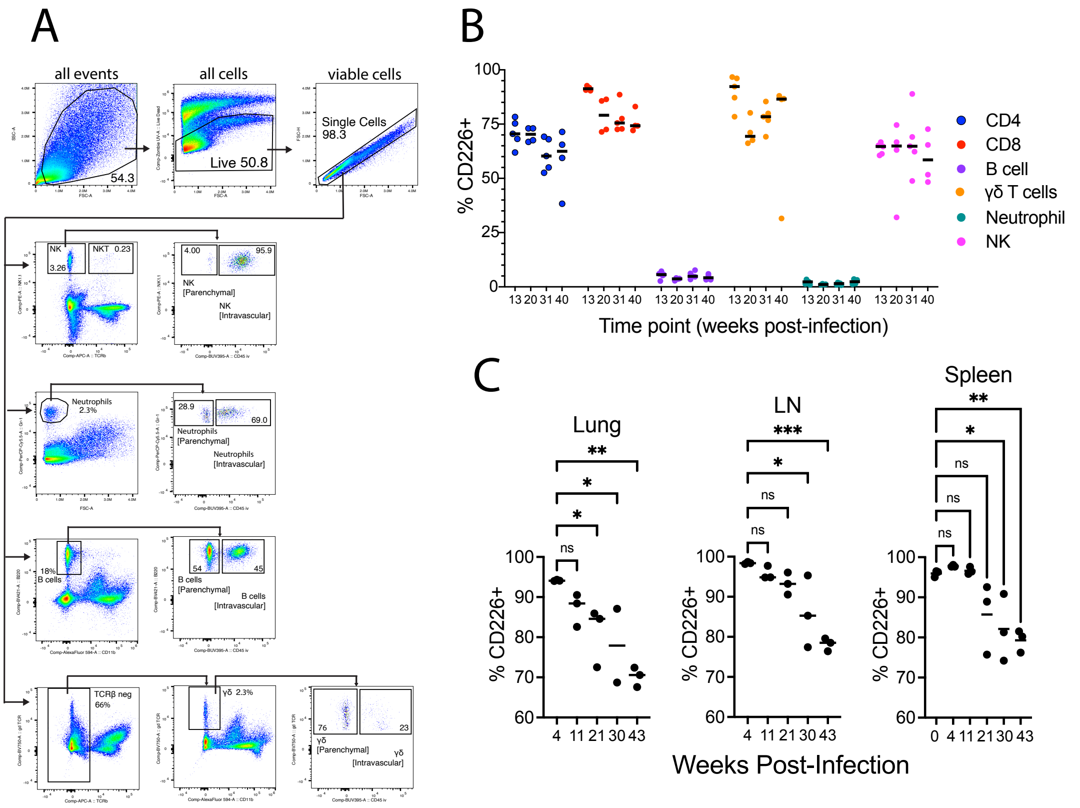


**Supplemental Figure 3. CD226 expression on lung immune cell subsets. (A)** Gating strategy for identifying lung immune cell subsets from mice infected with Mtb by flow cytometry. Numbers in the drawn gates are percentages. **(B)** Frequency of $CD226^+$ cells among lung parenchymal (CD45 $IV^-$) immune cells from mice infected with Mtb for indicated weeks. **(C)** Frequency of $CD226^+$ cells among antigen experienced ($CD62L^-CD44^+$) $CD8^+$ T cells in the lung parenchyma, mediastinal LN and spleen from mice infected with Mtb for indicated weeks. Splenocytes from uninfected mice are shown as 0 wpi. Data from n=4-5 mice/group (B) or 2-3 mice/group (C) and is representative of 2 (B) or 3 (C) experiments. Statistical testing used a One-way ANOVA and Tukey's multiple comparisons test. *, $p < 0.05$; **, $p < 0.01$; ***, $p < 0.005$; ns, not significant.

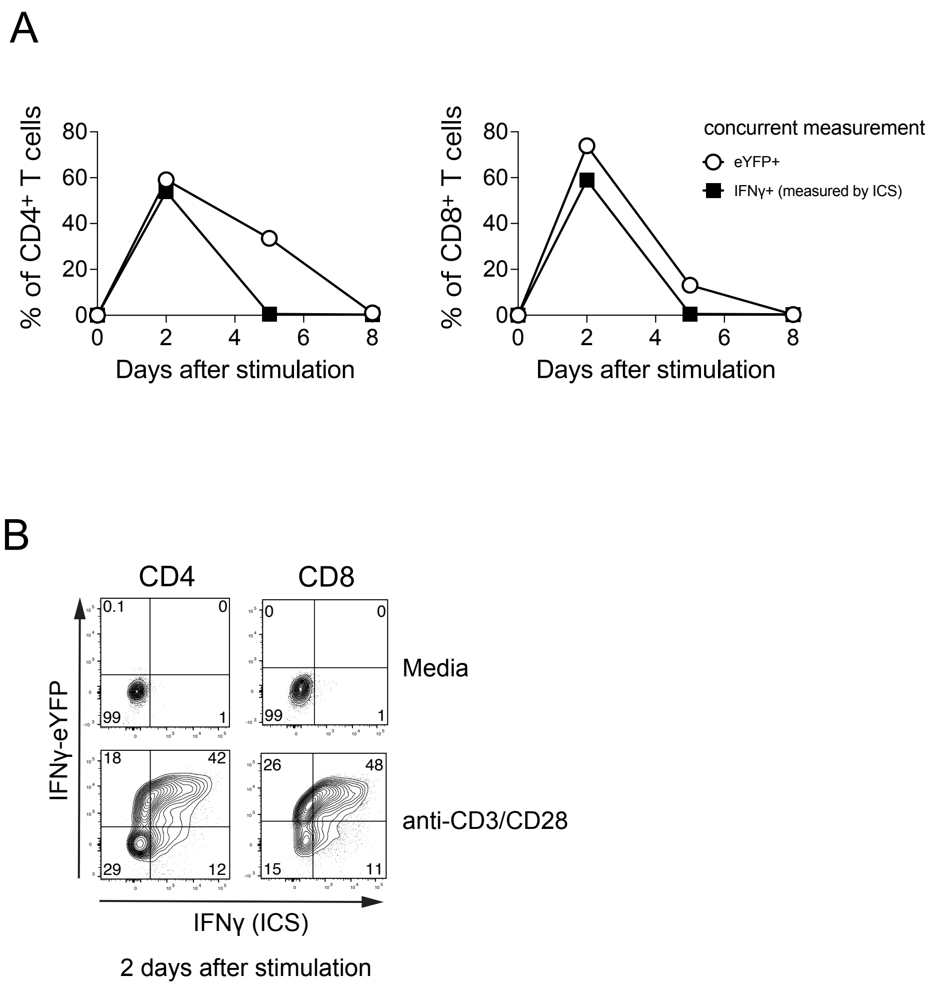


**Supplemental Figure 4. IFNγ-eYFP expression correlates with intracellular IFNγ measured by ICS. (A, B)** Graph depicting time-course of IFNγ-eYFP and intracellular IFNγ protein expression by splenocytes isolated from uninfected IFNγ-eYFP reporter after stimulation with anti-CD3 and anti-CD28. Quantification (A) and Representative flow cytometry plots of T cells two days after stimulation (B).

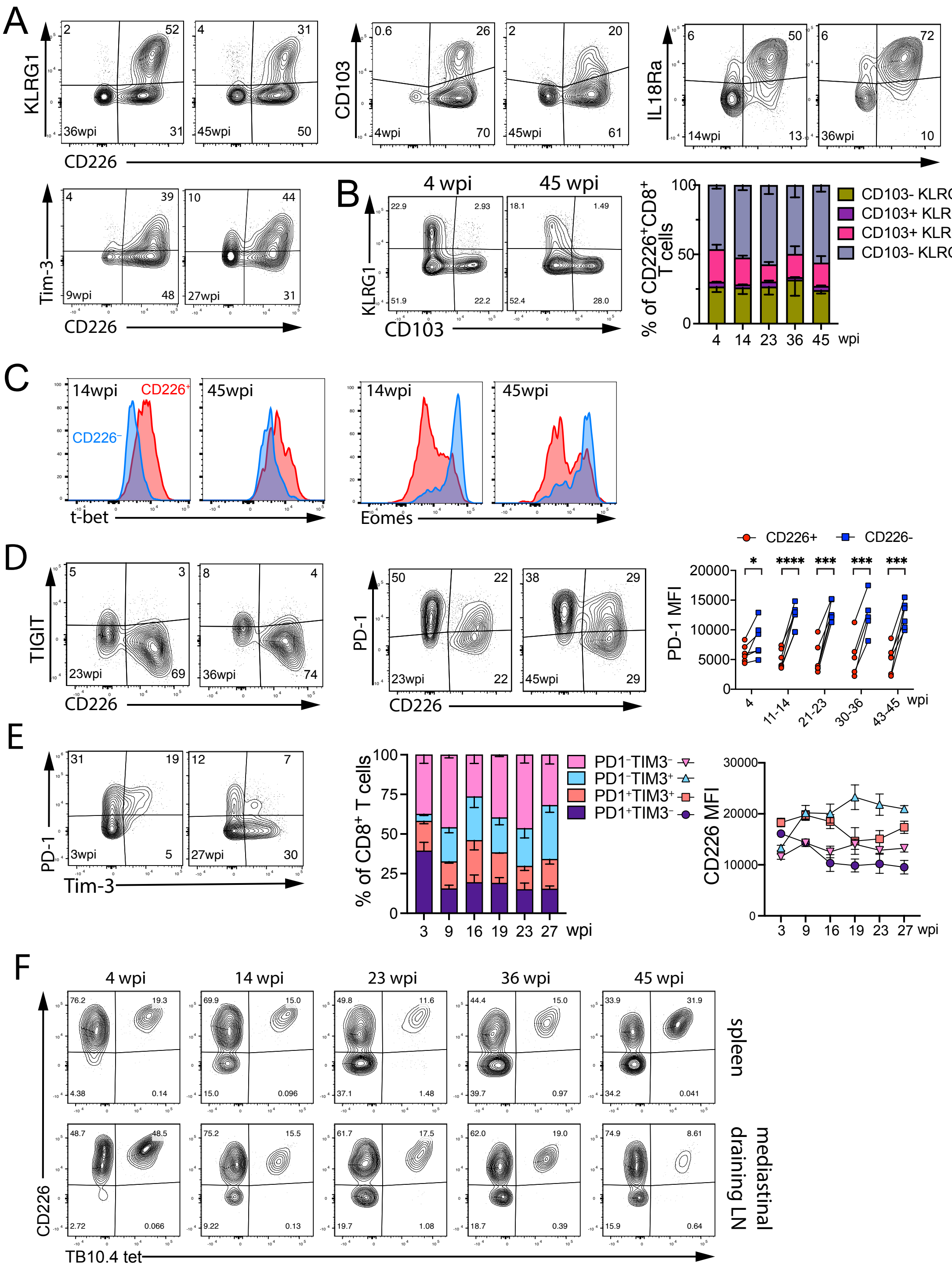


**Supplemental Figure 5. CD226 expression identifies terminally differentiated effector CD8+ T cells. (A)** Representative flow cytometry plots showing different phenotypic surface markers versus CD226 expression on lung parenchymal (CD45 IV⁻) antigen-experienced (CD62L⁻CD44⁺) CD8⁺ T cells from mice infected with Mtb for indicated weeks, related to Figure 5A, B. **(B)** Representative flow cytometry plots showing KLRG1 versus CD103 expression (left panel), and the bar plot depicting the proportions of subsets differentiated by CD103 and KLRG1 expression (right panel) on lung parenchymal (CD45-IV$^-$) antigen experienced (CD62L$^-$CD44$^+$) CD226$^+$ CD8$^+$ T cells from mice infected with Mtb for indicated weeks. **(C)** Representative flow cytometry histograms showing expression of TFs (T-bet and Eomes) on lung parenchymal (CD45 IV$^-$) antigen-experienced (CD62L$^-$CD44$^+$) CD8$^+$ T cells from mice infected with Mtb for indicated weeks, related to Figure 5C. **(D)** Representative flow cytometry plots showing TIGIT or PD-1 versus CD226 expression by lung parenchymal (CD45 IV$^-$) antigen experienced (CD62L$^-$CD44$^+$) CD8$^+$ T cells from mice infected with Mtb (left and middle panel), related to Figure 5D. Quantification of PD-1 MFI by lung parenchymal (CD45 IV$^-$) antigen experienced (CD62L$^-$CD44$^+$) CD226$^+$ and CD226$^-$ CD8+ T cells from mice infected with Mtb (right panel). **(E)** Representative flow cytometry plots showing PD-1 versus TIM-3 expression (left panel), the par plot depicting proportions of subsets differentiated by PD-1 and TIM-3 expression (middle panel) and CD226 MFI of indicated subsets (right panel) of lung parenchymal antigen experienced (CD62L$^-$CD44$^+$) CD8$^+$ T cells from mice infected with Mtb for indicated weeks. **(F)** Representative flow cytometry plots of CD226 expression by TB10.4$_{4-11}$-specific antigen experienced (CD62L$^-$CD44$^+$) CD8$^+$ T cells from spleen and mediastinal LN of mice infected with Mtb for indicated weeks. Numbers in the drawn gates are percentages. Data are representative of 2-3 experiments (2-3 mice/group). Data is mean ± SEM. Statistical testing used a paired, two-tailed Student's *t* test. *, $p < 0.05$; **, $p < 0.01$; ***, $p < 0.005$; ****$p < 0.001$; ns, not significant.

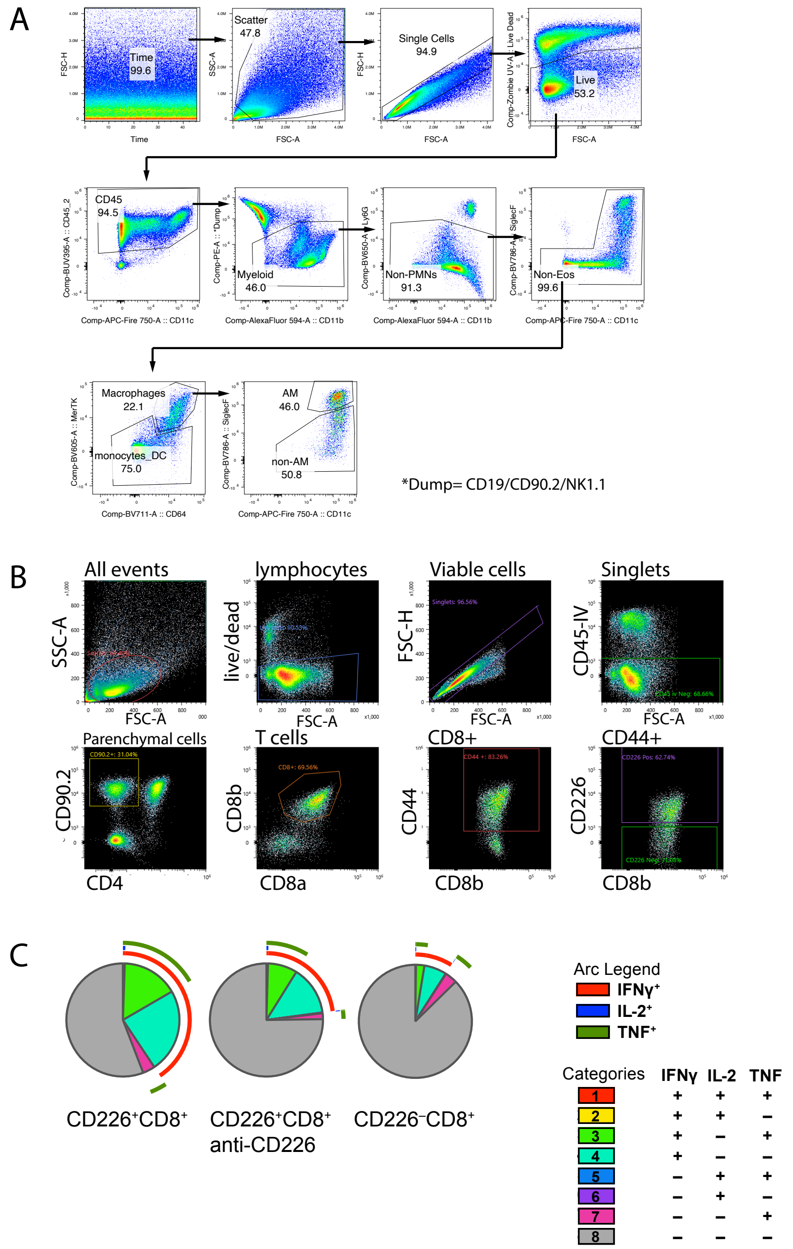


**Supplemental Figure 6. (A)** Gating strategy for identifying lung myeloid cell subsets from mice infected with Mtb by flow cytometry. PMN denotes polymorphonuclear leukocytes, Eos denotes eosinophils, DC denotes dendritic cells, and AM denotes alveolar macrophages. Numbers in the drawn gates are percentages. **(B)** Gating strategy for flow cytometric sorting of lung parenchymal $CD226^+$ and $CD226^-$ $CD44^+CD8^+$ T cells from mice infected with Mtb for in vitro stimulation. 3 mice were pooled for each sample, related to Figure 5E and Figure 6C, D, E**.** Numbers in the drawn gates are percentages. **(C)** Representative SPICE graph depicts the percentage of IFNγ, IL-2, and TNF expressing populations of sorted $CD226^+$ and $CD226^-$ $CD44^+CD8^+$ T cells from mice infected for 12 weeks stimulated with infected TG-PMs for 17 hours in the presence or absence of anti-CD226 antibody. 3 mice were pooled for each sample, related to Figure 6C, D, E.
